# Disturbance by soil mixing decreases microbial richness and supports homogenizing community assembly processes

**DOI:** 10.1101/2022.03.07.482922

**Authors:** Jaimie R. West, Thea Whitman

## Abstract

The spatial heterogeneity of soil’s microhabitats warrants the study of ecological patterns and community assembly processes in the context of community coalescence, or the combining and restructuring of communities and their environment. By mixing soil at various frequencies in a 16-week lab incubation, we explored the effects of mixing disturbance on soil bacterial richness, community composition, and community assembly processes. We hypothesized that well-mixed soil would harbor less richness, dominated by homogenizing dispersal and homogeneous selection. Using 16S rRNA gene sequencing, we inferred ecological processes, estimated richness and differential abundance, calculated compositional dissimilarity, and constructed co-occurrence networks. Findings supported our hypotheses, with >20% decrease in soil bacterial richness in well-mixed soil. While soil mixing resulted in increasingly dissimilar communities compared to unmixed soil (Bray-Curtis dissimilarity; 0.75 vs. 0.25), well-mixed soil communities were increasingly self-similar. Our results imply that vast soil diversity may be attributed to the unmixed and spatially heterogeneous nature of soil, and also provide insight into soil communities following coalescence events. By isolating and better understanding the effect of spatial heterogeneity and dysconnectivity on soil microbial communities, we may better extrapolate how anthropogenic disturbances, such as climate change or land use change, may affect broad soil functions.

**One sentence summary:** Soil mixing decreases bacterial richness as several taxa dominate the community, providing evidence for homogenizing community assembly processes.

## Introduction

Soil is a staggeringly complex, heterogeneous, and even harsh web of microhabitats that harbors vastly diverse communities of largely uncharacterized microorganisms that drive crucial soil functions such as biogeochemical cycling, organic matter decomposition, and plant productivity (Fierer 2017; Tecon and Or 2017). This diversity is potentially underpinned by the disconnected nature of soil microhabitats (Treves *et al.* 2003; Carson *et al.* 2010) and spatial soil heterogeneity (Fierer and Jackson 2006; Rillig, Muller and Lehmann 2017). Soil microhabitats range from largely undisturbed, as in deep soil horizons of a perennial grassland, to highly disturbed, as in the plough layer of an agricultural field. What are the features of the soil environment that support high levels of microbial diversity in soil? By reducing heterogeneity, changing resource availability, and forcing microbial interactions, how does soil disturbance change soil microbial diversity? It is essential to define and quantify the processes that generate and maintain microbial diversity in order to determine the mechanisms that drive soil functions and ecological relationships and to predict changes to soil biodiversity under global change.

Ascribing processes and mechanisms of community assembly to observed ecological patterns is a central question within ecology. Niche theory posits that community membership is governed by deterministic (non-random) processes driven by differences in the environment and fitness (Grinnell 1917; Hutchinson 1957; Chase and Leibold 2003, 2014). Alternatively, neutral theory asserts that community membership results from stochastic processes, or random changes in demographics and immigration acting on individuals and populations of equivalent fitness (Bell 2001; Hubbell 2001). Both theories accurately predict community patterns observed at various scales, and so the objective in a given system shifts towards determining the relative importance of niche and neutral processes, which together yield ecological assembly patterns (Adler, Hillerislambers and Levine 2007). Neutral theory holds power in its absoluteness, and operationally acts as a null hypothesis; where it fails, we may be able to identify deterministic processes (Gewin 2006). The mechanisms by which patterns and community membership unfold are generally categorized into four ecological processes: selection, drift, dispersal, and speciation (Vellend 2010; Zhou and Ning 2017). The importance and relative contributions of specific ecological processes for soil functioning are variable and poorly defined (Hubbell 2001; Hanson *et al.* 2012) and, as a result, ecological patterns are often weakly ascribed to the catch-all ‘black box’ of unknown aspects of community ecology (Vellend 2010).

The ecological processes of soil microbial community assembly that we focus on in this paper are dispersal, drift, and selection. Dispersal describes the movement and establishment of organisms in space, and is typically considered to be stochastic, as in passive dispersal via wind or water, and dependent upon population sizes. Dispersal may also be deterministic when driven by active, trait-dependent transport and establishment (Zhou and Ning 2017). Homogenizing dispersal results in higher compositional similarity between connected communities than would be expected by chance and stochasticity. Alternatively, dispersal limitation reflects higher compositional differences between communities than would be expected by chance, perhaps allowing for a stronger influence of drift—the stochastic changes in community membership attributable to reproduction and death (Stegen *et al.* 2013). Selection refers to deterministic or niche-based processes dictated by biotic factors such as inter-taxa fitness differences, and abiotic factors such as environmental filters (Hutchinson 1957). Homogeneous selection describes community assembly under similar conditions or filters, resulting in lower phylogenetic differences between communities than expected by chance (Dini-Andreote *et al.* 2015). Variable selection occurs when variable conditions produce different selective pressures, thus resulting in higher phylogenetic differences between communities than expected by chance (Stegen *et al.* 2015). Community assembly that occurs with only weak selection and stochastic dispersal is termed undominated, and encompasses ecological drift. Selection is perhaps the most intuitive of the community assembly processes, and can robustly shape and influence soil microbial communities, as in the global relationship between soil pH and bacterial richness and composition (Fierer and Jackson 2006; Fierer 2017). To statistically infer the relative influences of these community assembly processes in soil microbial communities, Stegen et al. (2012, 2013, 2015) have developed a null modeling approach that compares observed phylogenetic distance and dissimilarity metrics between communities to null models of stochastically assembled communities, originally demonstrated with river sediment communities (Stegen *et al.* 2012).

Despite robustly defined statistical models, characteristics of the soil environment and inhabitant microorganisms interact to result in relationships that can be difficult to predict *a priori*, in terms of their influence on community assembly processes (Evans, Martiny and Allison 2017). For instance, small populations of rare taxa, which constitute the majority of soil microbes (Lynch and Neufeld 2015), are more vulnerable to dispersal limitation and thus subject to drift, whereas larger populations of dominant soil taxa are typically governed by deterministic processes (Hanson *et al.* 2012). Even soil pH, arguably the strongest predictor of soil bacterial community composition (Delgado-Baquerizo *et al.* 2018), eludes consistent relationships to community assembly processes: selection can be strong under extreme pH conditions, but stochastic processes tend to characterize communities in soil with neutral pH (Tripathi *et al.* 2018). Soil aggregation, as a source of spatial heterogeneity, can shape microbial communities by furnishing distinct microhabitats (i.e. conditions for variable abiotic selection) or engendering microbial community isolation (i.e. inducing dispersal limitation) (Rillig, Muller and Lehmann 2017; Wilpiszeski *et al.* 2019), with individual aggregates potentially harboring unique communities (Bailey *et al.* 2013). Soil aggregate destruction or rearrangement may reduce spatial heterogeneity and give rise to homogenizing community assembly processes, such as homogenizing dispersal via movement of organisms, and homogenizing selection through more uniform allocation of resources and abiotic conditions. Because soil bacteria typically live in spatially-structured biofilms or microbial hotspots (Kuzyakov and Blagodatskaya 2015), disrupting soil structure would be expected to fundamentally alter community composition.

As scientists, we struggle to effectively study or even intuitively grasp the scale at which patchy communities of soil microbes function and interact with each other and their environments (Vos *et al.* 2013; O’Brien *et al.* 2016). Analyses of DNA extracted from a typical 250 mg soil sample often assume connectivity amongst the, perhaps, 300 million bacterial cells representing over 3000 operational taxonomic units (OTUs) that may be identified in a sample of that size, which can result in faulty ecological inferences (Rillig *et al.* 2015; Armitage and Jones 2019). In reality, the ‘sparsely populated, frequently dehydrated, maze’ of soil offers limited connectivity to its bacterial inhabitants, who are largely dependent upon moisture and biofilms for direct and indirect interaction (Or *et al.* 2007), and may only interact in small communities of perhaps 120 individuals (Raynaud and Nunan 2014). Soils are likely better thought of as vast collections of intermittently connected communities that often move and interact in units, such as in association with a soil particle or aggregate.

The movement and dispersal of microbes, along with mixing and restructuring of the spatially heterogeneous environments that they inhabit, has been termed community coalescence (Rillig *et al.* 2015, 2016), and this framework explicitly includes environmental mixing as a potential driver of community composition (Mansour *et al.* 2018). Coalescence events are relevant across many microbial environments, and frequently describe the convergence of disparate ecosystems with separate source communities (Rillig *et al.* 2015; Mansour *et al.* 2018), such as in aquatic systems where freshwater and brackish communities meet (Rocca *et al.* 2020), the human microbiome as microbes in food are ingested (Pasolli *et al.* 2020), industrial anaerobic digestors where waste is seeded with degrading communities (Sierocinski *et al.* 2017), and in soil management where a consortium of microbes with specific functional traits are added to restore degraded soil (Calderón *et al.* 2017). Community coalescence in soil has particular nuances. Soil mixing events are likely to be spatially fragmented, leaving much of the soil relatively undisturbed, as compared to aquatic systems, which are more readily homogenized (Philippot, Griffiths and Langenheder 2021). This is one possible mechanism by which soil maintains such high levels of diversity (due to variably affected sub-communities) and functional resilience (maintained within undisturbed communities) (König *et al.* 2019). In addition to describing the mixing of distinct environments, coalescence is also applicable to any physical disturbance of soil, where there is a high level of inherent heterogeneity in seemingly uniform environments. For instance, coalescence occurs when an earthworm’s drilosphere meets the rhizosphere (Jacquiod *et al.* 2020), through agricultural tillage (Guillou *et al.* 2019) and during cryoturbation (Gittel *et al.* 2014).

We sought to explore the effects of community coalescence in soil—What happens to microbial diversity and community assembly processes when soil is mixed to coalesce isolated communities and heterogeneous microhabitats? We hypothesized that well-mixed soil would harbor a less diverse microbial community, and we predicted that community coalescence in soil would decrease richness and result in increasingly homogeneous communities dominated by homogenizing dispersal and homogeneous selection. Our goals were to relate differences in richness, compositional (dis)similarity, and relative contributions of community assembly processes—namely dispersal and selection—to soil mixing. To address these goals, we subjected the soil environment to mixing at various frequencies and assessed the outcomes on soil microbial communities and associated community assembly processes. Previous work has investigated differences in ecological processes across habitats (Stegen *et al.* 2012; Huber *et al.* 2020), for which selection via environmental filtering would be expected to be the dominant process (Hanson *et al.* 2012; Tripathi *et al.* 2018); we sought to shift our focus from the predominant influence of selection that is inherent across environmental gradients and, rather, focus on the potential for selection and dispersal as it pertains to the spatially-structured soil environment by manipulating a single soil in a lab setting. Complementary objectives of this work were to explore community composition, and, to test our methodological approach, in which we attempted to balance the microscale at which microbes interact with each other and the soil environment, and the smallest scale at which soil and soil conditions can reasonably be manipulated in the lab without resorting to simulated soil environments or robotic handling systems.

Here, we characterize the microbial communities that inhabit about 50 mg of soil (approximately the size of a lentil) following a 16-week incubation experiment in which we coalesced the soil environment and its inhabitant soil communities by combining, mixing and re-dividing soil “sets” at various frequencies. We used 16S rRNA gene sequencing to determine community membership, and evaluated community assembly processes using statistical models. This work quantifies community coalescence in soil, thus providing insight on the effects of coalescence events, such as agricultural tillage or bioturbation. By isolating and better understanding the effects that abiotic factors and spatial heterogeneity can have on community assembly processes in soil, we are also better equipped to extrapolate the effects that anthropogenic processes, such as climate change or land use change, may have on broad soil functions.

## Methods

### Soil collection

Our priorities in collecting soil for lab manipulation were to: (1) obtain a fairly diverse initial soil community to assess community changes; (2) obtain an unmanaged soil, to minimize effect of management or prior amendments; and (3) obtain a soil with moderate or low clay content, to minimize extracellular DNA (Morrissey *et al.* 2015) and mitigate potential for soil compaction and stickiness during manipulation. As such, we collected Freeon silt loam soil (a very deep, moderately well drained, coarse-loamy, mixed, superactive, frigid Oxyaquic Glossudalf; Luvisols by WRB classification) on 30 August 2018 near Connor’s Lake in Sawyer County, Northern WI, U.S. (45°44’55.6”N, 90°43’47.1”W, 430 m asl) (SI Fig. 1). Vegetation type was northern mesic forest, early-to-mid seral, dominated by *Acer rubrum* L. (approx. 80%), *Acer saccharum* Marsh. (approx. 10%), *Betula alleghaniensis* Britt. (approx. 5%), and *Tilia americana* L. (<5%). Two soil cores (1.8 cm diameter) were collected from each of six locations randomly chosen along a 50 m transect. From each of these 12 soil cores, we retained a portion of the A horizon, 15–20 cm of depth, in a large Whirl-Pak bag kept on ice prior to refrigeration. A representative subsample of air-dried soil was submitted for standard analyses and was found to be comprised of 50% silt, 36% sand and 14% clay (hydrometer method) (Bouyoucos 1962); 2.7% organic matter (loss on ignition) (Schulte and Hopkins 1996), 5.1 pH (1:1 water) (Richards, 1954), 13 mg P kg^−1^, 22 mg K kg^−1^ (P and K by Bray-1 method) (Bray and Kurtz, 1945), 172 mg Ca kg^−1^, 25 mg Mg kg^−1^ (Ca and Mg by ammonium acetate method) (Thomas 1982) and < 3 mg available N kg^−1^ (NO_3_^−^–N + NH_4_^+^–N by KCl extraction) (Doane and Horwath 2003).

### Experimental setup and design

Briefly, to investigate the effects of mixing and community coalescence on soil microbial community ecological processes and diversity, we incubated sets of soils that, at various frequencies, were combined and mixed together by vortex, in addition to controls that were never combined or mixed after establishment, and vortex controls that were never combined with other soil. To establish the experiment, freshly collected, field-moist soil was gently shaken through an ethanol-sterilized sieve to 2 mm and homogenized, removing any visible roots. We then established eight-tube mixing sets, with each tube (0.5 mL freestanding tube, catalog no. 16466-036, VWR) containing 50 mg soil (± 5 mg), and each mixing set totaling 400 mg soil (SI Fig. 3). These eight-tube sets were randomly assigned to mixing treatments, which determined how frequently the soil in the set would be combined in one tube (2 mL freestanding tube, catalog no. 89004-308, VWR), mixed by vortex, and re-divided over a 16-week incubation period (Fig. 1). The mixing treatments included: two times mixed (2×; soil was manipulated at the beginning of the incubation and again halfway through the incubation), four times mixed (4×; every fourth week), eight times mixed (8×; every other week), 16 times mixed (16×; weekly), 32 times mixed (32×; twice weekly) (Fig. 1). There were four replicate mixing sets for each treatment. As a control, there were 32 tubes of soil, 50 mg (± 5 mg), that were incubated undisturbed for the duration of the experiment (1×, or control).

**Figure 1.**
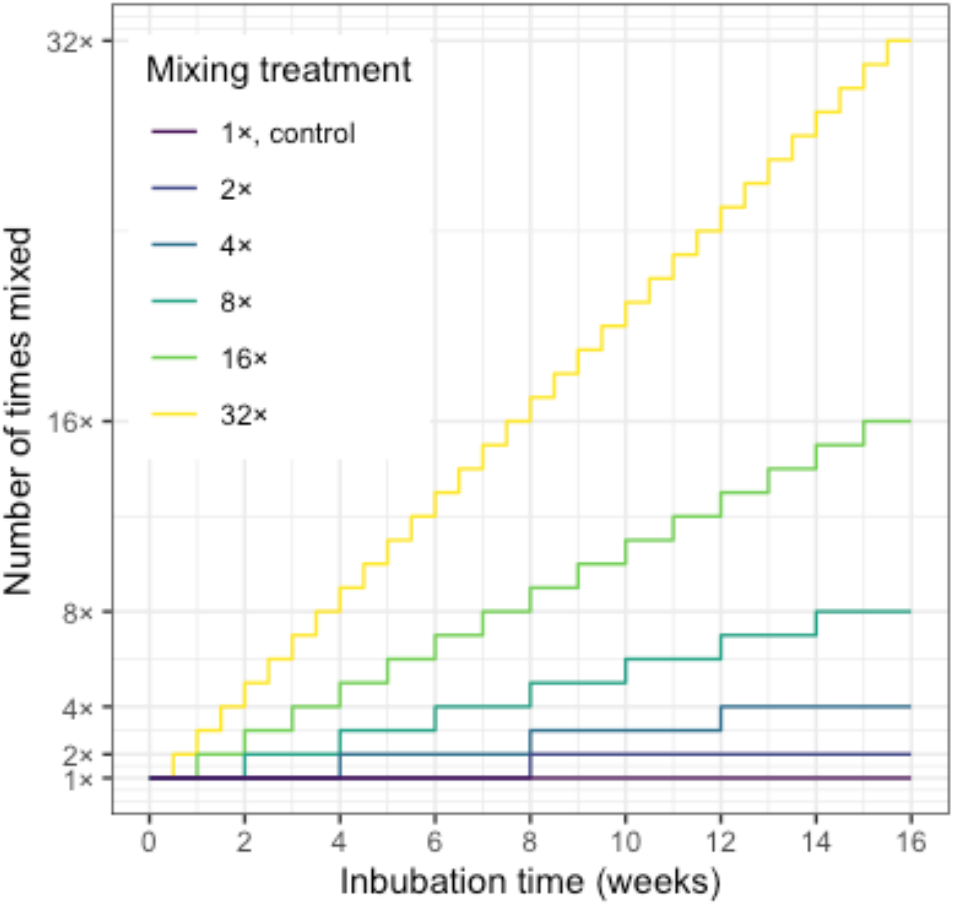
Visual representation of mixing treatments over the course of the 16-week soil incubation. The mixing treatment represents the number of times each mixing set was combined, mixed by vortex, and re-divided. The 1× controls (n = 32) were incubated for 16 weeks undisturbed following mixing at the time the experiment was established.

At respective times of mixing, the eight tubes within a given mixing set were weighed to gravimetrically measure moisture loss, and all soil within the mixing set was combined in one separate tube, with great care taken to transfer all soil and residue. Autoclaved Milli-Q water was added to the soil at a rate matching total moisture loss, and the tube was agitated using a vortex mixer (catalog no. 02215365, Fisher Scientific) fitted with a horizontal tube holder at speed seven for five seconds. Following vortex mixing, the soil was weighed and evenly divided back into the eight original incubation tubes (Fig. 2). To account for any effects of vortex mixing, there were concurrent vortex controls, which were stand-alone tubes that underwent moisture correction and vortex mixing, but were never combined with any other soil (Fig. 2). There were eight vortex control tubes per treatment (two associated with each mixing set) except the 1× control, which did not undergo subsequent mixing and therefore did not warrant vortex controls.

**Figure 2.**
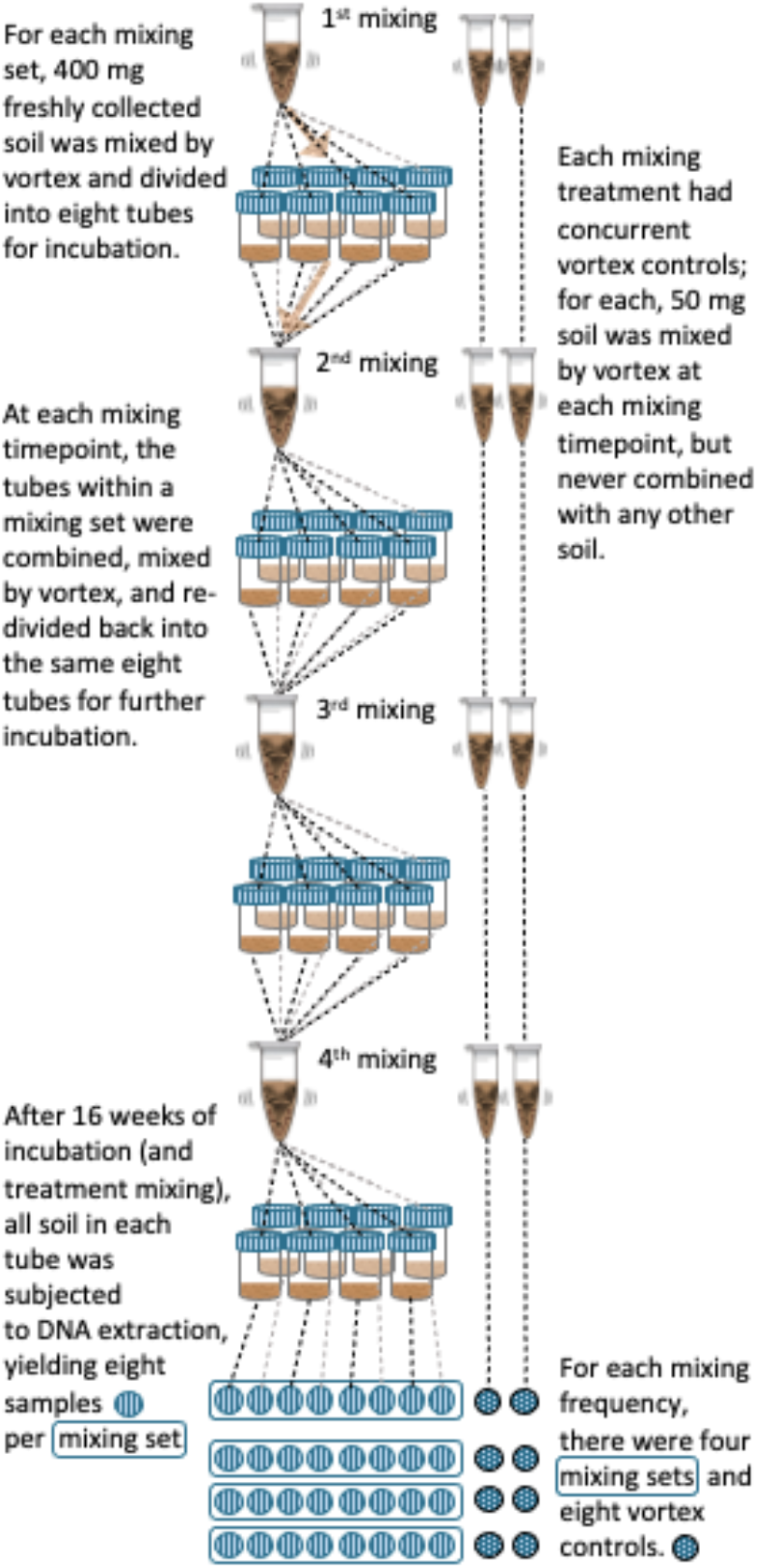
Experimental setup depicting one mixing set in the 4× mixing treatment, in which soil is combined, mixed by vortex, and re-divided at regular intervals. Note: the incubation interval varied in length depending on the mixing frequency. Concurrent vortex controls were stand-alone tubes of soil that were mixed by vortex at each mixing timepoint but never combined with other soil. At the conclusion of the 16-week incubation, DNA was extracted from soil in each incubation tube for 16S rRNA gene amplicon sequencing. See SI Fig. 2 for an expanded version of this figure that includes all treatments.

The cap of each incubation tube had one 1/32” hole for air exchange, drilled at a 45° angle (for the vortex controls, a clean cap without an air hole was briefly used during vortex mixing). All tubes and caps were autoclaved prior to use. Tubes were incubated in two identical dark incubation boxes at room temperature and > 95% relative humidity (RH) to reduce soil drying (SI Fig. 3). The RH was maintained with open Petri dishes containing autoclaved MilliQ water. Temperature and RH were continuously monitored in each incubation box (data not shown). The incubation boxes were frequently opened for treatment manipulation, and thus kept aerated.

In order to characterize the microbial community at the time of soil sampling, we also retained 32 x 50 mg (± 5 mg) soil samples, which were frozen at −80 °C without incubation (“Initial”). At the conclusion of the experiment, all tubes were frozen at −80 °C prior to DNA extraction. An electrode deionizer (catalog no. 05.8091.100, Haug North America, Mississauga, ON, Canada) and antistatic nitrile gloves were used while manipulating the soil to minimize static attraction and repulsion.

### DNA extraction and 16S rRNA gene sequencing

Total genomic DNA was extracted from all soil within each incubation tube, which ranged from 30–55 mg soil per tube at the end of the experiment. Care was taken to transfer all soil residue and DNA through a series of washes with PowerBead Solution in conjunction with vortex agitation. Complete library preparation details can be found in the Supplementary Information. Briefly, the 16S rRNA genes of extracted DNA were amplified in triplicate using PCR. Variable region V4 of the 16S rRNA gene was targeted using forward primer 515f and reverse primer 806r (Walters *et al.* 2016). Amplified DNA was normalized and purified, prior to paired-end 250 base pair sequencing on an Illumina MiSeq sequencer at the UW-Madison Biotech Center. To obtain high coverage, the same library was sequenced twice under identical conditions, and total reads were pooled for each sample after processing as described next. Sequencing data was processed using a QIIME2 (Bolyen *et al.* 2019) pipeline, with DADA2 (Callahan *et al.* 2016) as the OTU (or amplicon sequence variant)-picking algorithm, and taxonomy assignment using the SILVA 132 reference database (Quast *et al.* 2013; Yilmaz *et al.* 2013). This yielded 16 687 633 demultiplexed sequences, which was reduced to 12 961 153 after denoising, with a mean length of 238 base pairs (SD = 5.5). Excluding blanks, a total of 9264 OTUs were identified. Amplicon sequences will be made available in the National Center for Biotechnology Information (NCBI) Sequence Read Archive (SRA) upon publication. Our primers targeted both bacteria and archaea, but because our communities were dominated by bacteria (> 99.2% of total reads), for simplicity, we will refer to bacteria when discussing communities in this manuscript, but recognize this includes archaea, with over 96% of archaeal reads representing the phylum *Crenarchaeota*.

### Community assembly process assignments

To identify the community assembly processes associated with soil mixing and community coalescence, we adapted a null-modelling method (Stegen *et al.* 2012, 2013, 2015) (R code adapted from https://github.com/stegen/Stegen_etal_ISME_2013). This approach used the abundance-weighted beta-mean nearest taxon distance (βMNTD; the mean phylogenetic distance between each OTU in one community and its closest relative in another community) (Fine and Kembel 2011), and Bray-Curtis dissimilarities (BC-Dis) for pairs of communities (Bray and Curtis 1957). These values were then compared to null distributions of equivalent values in order to determine whether there was a significant influence of selection and/or dispersal, respectively. Unlike applications of this model to different field-based communities, which require a series of assumptions to calculate a null distribution, we had a “true” baseline or null scenario in the 1× control soil condition. Values from all mixing treatments were thus compared to values from these 1× communities to determine the relative effects of selection and dispersal. The 32 1× control samples were incubated alongside the mixing treatments, undisturbed after the initial soil homogenization and mixing, and thus represented stochastic community assembly in absence of mixing-induced selection or dispersal pressure. We created the null distributions for both βMNTD and BC-Dis by randomly grouping the 1× controls into four sets (eight tubes each) and calculating these metrics for each pair of tubes within a randomly grouped eight-tube set. This allowed us to ask the question, “Did a given mixing treatment increase the influence of any community assembly processes as compared to unmixed communities?” Detailed methods for community assembly processes assignment follow.

To determine the influence of selection, the null distribution values of βMNTD (βMNTD_Null_) were arranged in ascending order and the 95% confidence interval (CI) was nonparametrically identified by finding the 0.025 and 0.975 quantiles. We then took the observed βMNTD (βMNTD_Obs_) values for every possible pair of communities within a mixing set and compared that to βMNTD_Null_. Any comparison for which βMNTD_Obs_ was below the 95% CI of βMNTD_Null_ was assigned to **homogeneous selection**. Comparisons that fell above the 95% CI were assigned to **variable selection**. Comparisons that fell within the 95% CI of βMNTD_Null_ values were considered to lack a strong influence of selection.

To determine the influence of dispersal, the null distribution values of BC-Dis (BC-Dis_Null_) were arranged in ascending order and the 95% CI was nonparametrically identified by finding the 0.025 and 0.975 quantiles. We then took the observed BC-Dis (BC-Dis_Obs_) values for every possible pair of communities within a mixing set and compared them to the BC-Dis_Null_. Any comparisons that were below the 95% CI of BC-DisNull values were assigned to **homogenizing dispersal**, and comparisons above the 95% CI were assigned to **dispersal limitation**. Comparisons with BC-Dis_Obs_ values that fell within the 95% CI for BC-Dis_Null_ values were considered to reflect stochastic dispersal. Some comparisons fell below the 95% CI in both measures (**homogeneous selection & homogenizing dispersal)**, or above the 95% CI in both measures (**dispersal limitation & variable selection)**. Community pair comparisons that fell within the 95% confidence interval for both metrics were considered **undominated**by any particular community assembly process because compositional differences exhibited weak selection and random dispersal. Ecological drift may drive large differences in communities under dispersal limitation, and may be important in undominated communities, where selection and dispersal are weak (Vellend 2010). The potential for dual assignment of both selection and dispersal is one way in which our approach differs from that of Stegen *et al.* (2012, 2013, 2015), which precludes identification of dispersal if there is evidence for selection. This decision hinges on the premise that selection, when present, is expected to overpower the effects of dispersal processes. However, these processes are not mutually exclusive, and the metrics described above can be used together to capture instances when selection and dispersal (i.e. deterministic and stochastic processes) could simultaneously be influencing community assembly. Other recent approaches have developed models to assign community assembly processes separately to different ‘bins’ of OTUs (Ning *et al.* 2020; Fodelianakis *et al.* 2021), though these methods risk overstating relationships between community processes and perhaps idiosyncratic OTU bins.

This entire process (randomly assigning 1× controls into sets, calculating βMNTD_Null_ and BC- Dis_Null_, establishing 95% CI cutoffs for each null iteration, and assigning community assembly processes in mixing treatments relative to the null) was iterated 999 times. The range and distribution of observed values and 95% CI cutoffs are visualized in SI Fig. 4. Anecdotally, as few as 10 iterations yielded very similar results for this dataset, making this a computationally tractable test.

### Data analysis

Data analysis was performed in R (R-Core-Team 2018), version 4.0.3, using *ggplot2* (Wickham 2016) for data visualization. To describe diversity, we used the weighted linear regression model of OTU richness estimates, which weights observations based on variance, to calculate 95% CIs for treatment means using the *betta()* function in *breakaway* for R (*breakaway∷betta)* (Willis and Bunge 2014), interpreting only treatments with non-overlapping 95% CIs. Beta diversity was visualized for Bray-Curtis dissimilarities (Bray and Curtis 1957) of relative abundance data using principal coordinates analysis (PCoA) created with *phyloseq∷ordinate* (McMurdie and Holmes 2013). To test for a significant effect of mixing treatment on community composition, we used permutational multivariate analysis of variance (PERMANOVA) to partition Bray-Curtis dissimilarity matrices among sources of variation using *vegan∷adonis* (Anderson 2001). A significant result (p < 0.05) was subjected to *post-hoc* pairwise comparisons, adjusting p-values using the Benjamini-Hochberg method (Benjamini and Hochberg 1995) to identify significant differences between mixing treatments. To test if treatments differed in their dispersion, we used homogeneity of multivariate dispersions (PERMDISP; *vegan∷betadisper*), a resemblance-based permutation test of the null hypothesis that the within-group dispersion is equivalent among groups; non-homogeneous dispersion of the data would result in rejection of the null hypothesis (Anderson 2006). To quantify the degree to which treatments differed from the 1× control, as well as how much tubes differed within mixing sets, we used Bray-Curtis dissimilarities (*vegan∷vegdist*) (Oksanen *et al.* 2019) and ANOVA to test for a significant treatment effect. A significant result (p <0.05) was subjected to *post-hoc* multiple comparisons to 1× controls using Dunnett’s Test (Dunnett 1955). When assigning community assembly processes as described above, selection was inferred by βMNTD (*picante∷comdistnt*) (Kembel *et al.* 2010). Dispersal was inferred as described above by calculating Bray–Curtis dissimilarities (*phyloseq∷distance*) on OTU relative abundances.

After evaluating our key questions, we assessed differential abundance to identify significant treatment-driven shifts in relative abundances of taxa. For this analysis, we compared each treatment to the 1× control (excluding taxa with mean relative abundance < 0.00001) and subjected these datasets to a beta-binomial regression model and “Wald” hypothesis test in *corncob∷differentialTest* (Martin, Witten and Willis 2021), which controls for the effect of the mixing treatment on dispersion. We report the μ value, which is the coefficient used to estimate relative abundance in the *corncob* model and is proportional to the fold-change in relative abundance between the treatment and control. To further understand changes in community composition, we sought to test the relationship between mixing treatment and mean predicted rRNA gene copy number, which may correlate with potential growth rate, by calculating the weighted mean predicted 16S rRNA gene copy number for each sample (Nemergut *et al.* 2016) and compared treatments using ANOVA and *post-hoc* testing, as described above. Predicted rRNA gene copy numbers were obtained using the ribosomal RNA operon database (rrnDB) (Stoddard *et al.* 2015). To determine co-occurrence of bacterial OTUs across samples within the same treatment, we used network analysis (Connor, Barberán and Clauset 2017) following the approach of (Whitman *et al.* 2019), and visualization using *igraph* (Csardi and Nepusz 2006). After dropping small and disconnected network components to obtain the primary network, we identified modules of primarily co-occurring OTUs using edge correlation values to weight co-occurrences positively and co-exclusions negatively (*igraph∷cluster_spinglass*). For networks where the secondary network component was similar in size to the primary component, the secondary component was also analyzed. The R code used to perform these analyses and to create the following figures is available at https://github.com/jaimiewest/Soil-Mixing.

## Results

### Soil mixing decreased bacterial richness

Increased mixing frequency decreased bacterial richness (Fig. 3), with the most frequently mixed soil treatments (16×, and 32×) demonstrating lower richness than the 1× controls, 2×, 4×, and 8× treatments, as well as the initial soil. Notably, even as rate of vortex mixing increased, the vortex controls maintained a consistent level of richness not statistically different from that of the 1× controls.

**Figure 3.**
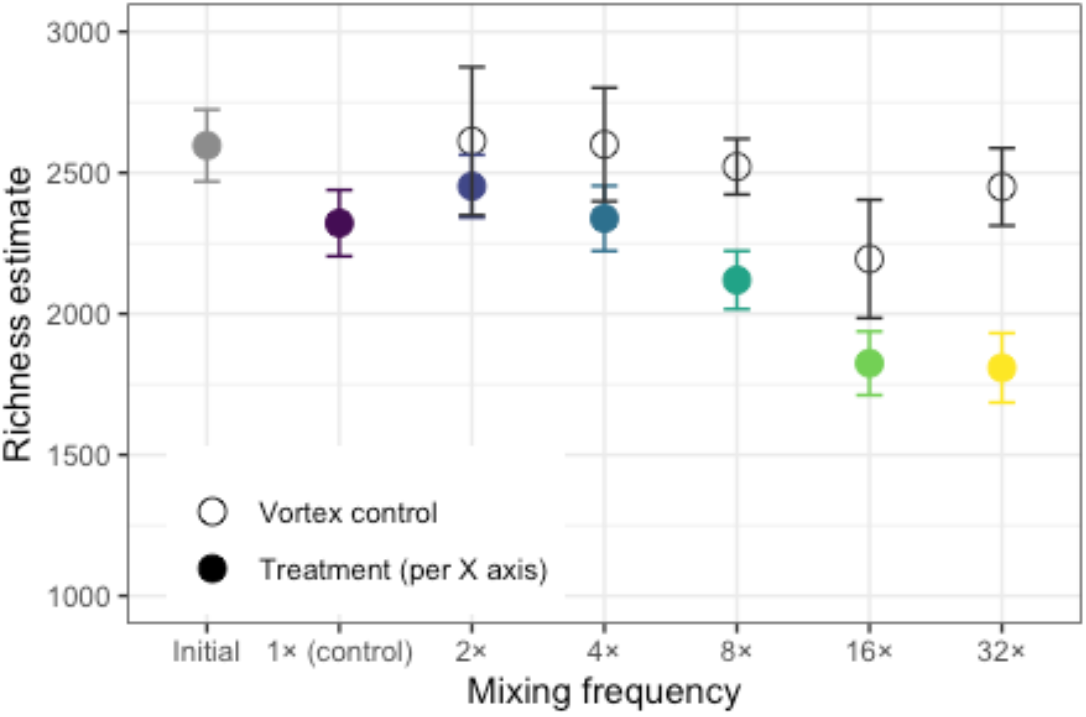
Community-level OTU richness, by mixing treatment (closed points), or frequency of vortex mixing (open points). Richness of the initial soil communities represents freshly collected soil that did not undergo incubation (gray point). Error bars represent 95% confidence intervals (± 1.96 * SE).

### Mixed soil communities became more similar to each other while diverging from unmixed controls

Mixed soil communities were significantly different from the unmixed 1× controls, as is apparent in both the ordination (Fig. 4; PERMANOVA, *p* = 0.001, R^2^ = 0.73, and *p* ≤ 0.001 for each pairwise comparison between mixing treatments) and in community dissimilarity (Fig. 5a; ANOVA, *p* < 0.0001, and significant pairwise comparisons, *p* < 0.05 for 2× and *p* < 0.0001 for all other treatments, Dunnett’s). Axis 1 of the PCoA (Fig. 4) explains 58% of the variation amongst these communities, which is the axis along which the mixing treatment spreads, with tubes of the same mixing treatments generally grouping together. Though mixed soil increasingly differentiated from unmixed soil, communities *within* each mixing set became more similar to each other with mixing (Fig. 5b; ANOVA, *p* < 0.0001, and *p* < 0.0001 for each pairwise treatment comparison to 1×, Dunnett’s), as is underscored by clustering within mixing sets in the PCoA, particularly within the 16× and 32× treatments (Fig. 4). The treatment-driven clustering pattern apparent in the PCoA ordination illustrates the importance of mixing frequency on soil microbial community composition data, though the results of the PERMANOVA may be affected by non-homogeneous dispersion in the data (*p* = 0.01, PERMDISP), e.g. the group of 1× control points are more tightly clustered, indicating lower dispersal of points within this treatment, compared to dispersal of points within mixed treatments (Fig. 4). The initial, unincubated samples were included in the PCoA in order to gauge the overall effect of the lab incubation on soil communities, which is much smaller than the effects of mixing.

**Figure 4.**
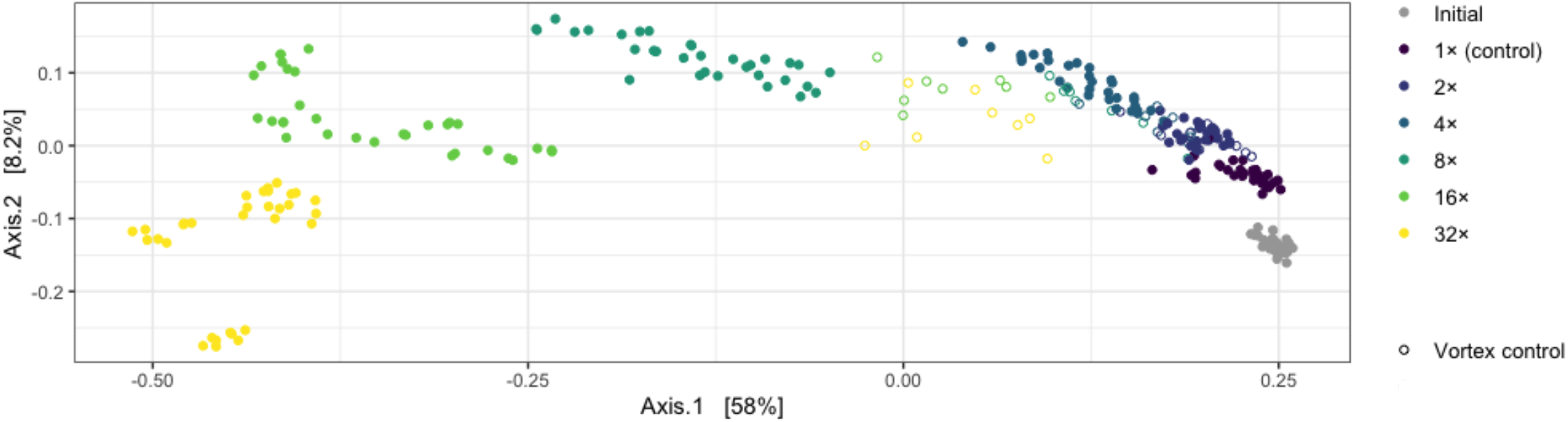
Principal coordinates analysis of Bray-Curtis dissimilarities of community data as relative abundance, colored by mixing treatment. Each point represents one tube (up to ~50 mg soil). Vortex controls (open points) were mixed by vortex but never combined with other soil. Initial communities (grey points) represent the community present in freshly collected soil that did not undergo incubation.

**Figure 5.**
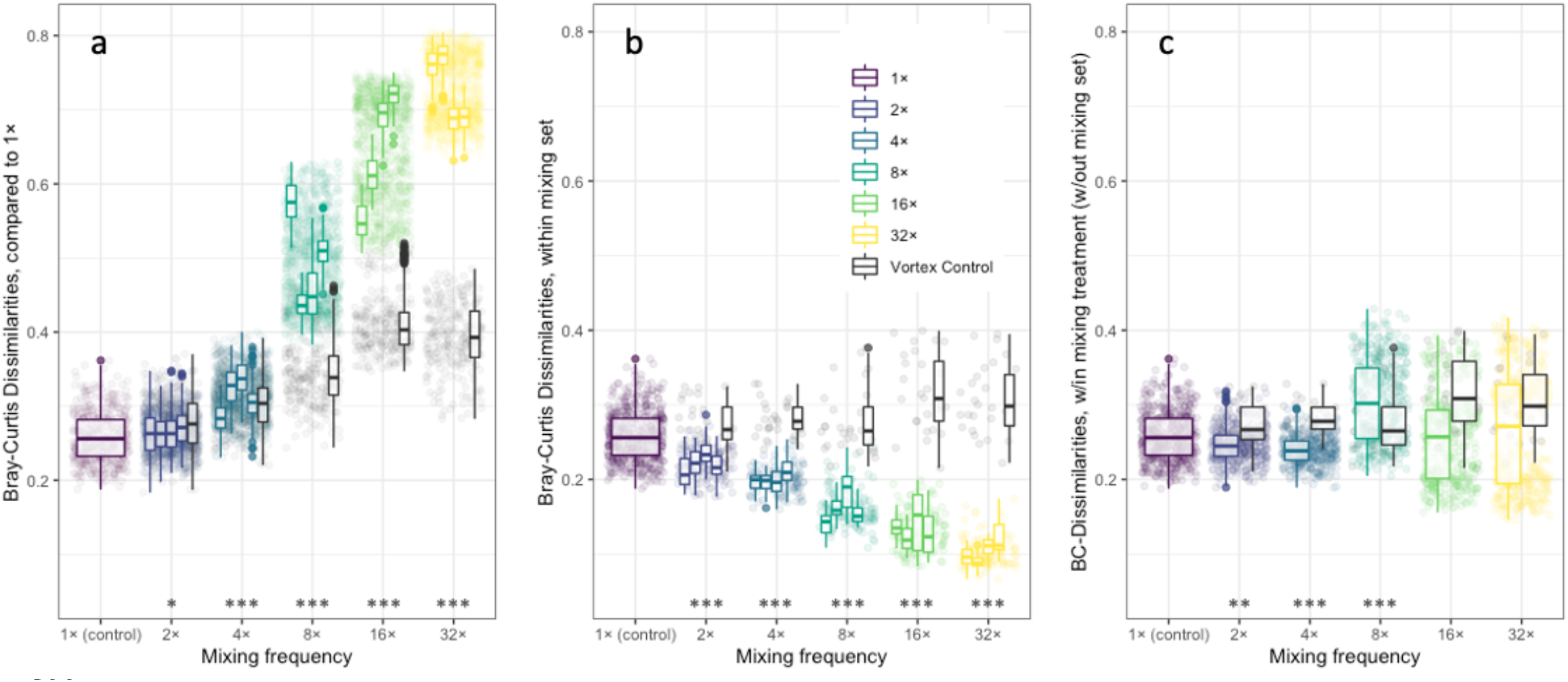
Bray-Curtis dissimilarities of bacterial community composition. (**a**) Each boxplot represents a mixing set and shows the dissimilarity compared to 1× controls, quantified for each possible pairwise comparison. There were four mixing sets per treatment, 2×–32×, and each mixing set had eight tubes. Additionally, there were eight vortex controls associated with each mixing treatment, which were stand-alone tubes mixed by vortex concurrently with mixing sets but never combined with other soil. There were 32 1× controls; the 1× boxplot shows pairwise comparisons amongst all 1× controls. (**b**) Each boxplot represents the dissimilarity within a mixing set for treatments 2×–32×, or amongst all 32 1× controls, which were not mixed during the incubation and thus had no mixing sets. (**c**) Each boxplot shows the dissimilarity amongst all tubes within the same soil mixing treatment, but excluding pairs of tubes in the same mixing set. Asterisks represent statistically significant treatment differences from 1×; *** = *p* < 0.001, ** = *p* < 0.01, * = *p* < 0.05, Dunnett’s test (statistics for vortex controls are reported in the text).

Vortex control communities were also significantly different in composition from the 1× controls, though to a lesser magnitude than the frequently mixed soil treatments (Fig. 5a in black; *p* < 0.0001, ANOVA; each treatment significant compared to 1×, *p* < 0.0001, Dunnett’s). However, vortex controls did not become more similar to each other with mixing; pairwise dissimilarities were significantly higher than those amongst 1× controls (Fig. 5b in black; *p* < 0.0001, ANOVA; *p* < 0.05 for 4× and 8×, *p* < 0.0001 for 16× and 32×, Dunnett’s).

To determine if there was an overall effect of the mixing treatment on community (dis)similarity, we compared communities that underwent the same mixing but were not mixed together, i.e. tubes from the same treatment but excluding tube pairs from the same mixing set (Fig. 5c). Compared to dissimilarity amongst 1× tube communities, we found a significant treatment effect (*p* < 0.0001, ANOVA), with significant decreases in Bray-Curtis dissimilarities at 2× and 4× (*p* < 0.01, Dunnett’s), and a significant increase at 8× (*p* < 0.0001, Dunnett’s). Dissimilarity amongst communities at 16× and 32× was not significantly different than dissimilarity amongst 1× tube communities (Fig. 5c), which stands in contrast to within-mixing set comparisons, in which communities became more similar with soil mixing (Fig. 5b).

### Community Assembly

Soil mixing altered the relative contributions of ecological community assembly processes (Fig. 6). Increased mixing frequency was marked by homogenizing community assembly processes; both homogenizing dispersal and homogenizing selection characterized nearly 100% of all pairwise comparisons at 16× and 32× (Fig. 6a). With less frequent soil mixing, there was a greater proportion of comparisons that were considered undominated by any particular process, with 27% and 72% undominated at 4× and 2×, respectively.

**Figure 6.**
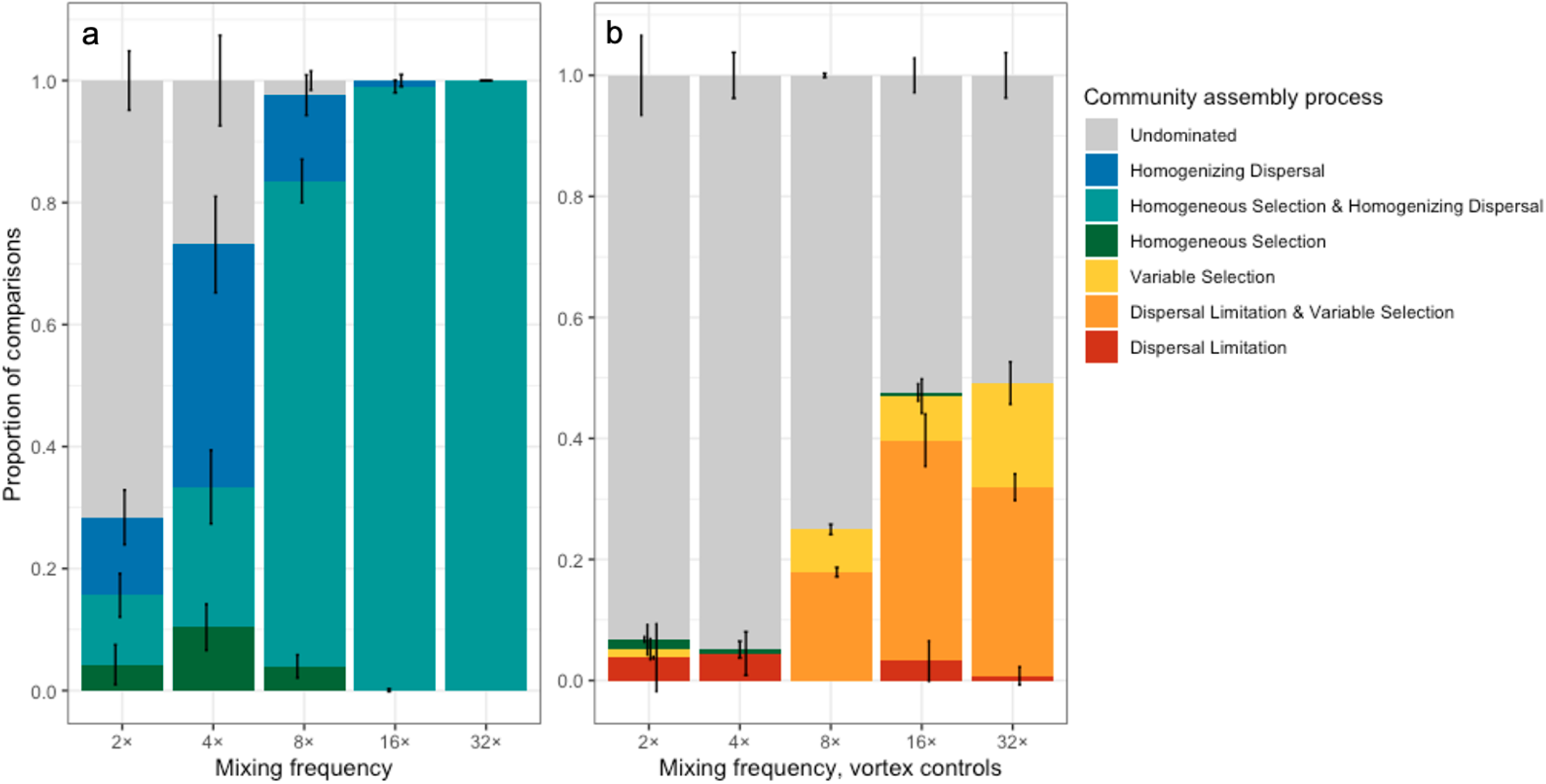
The community assembly process assignments as driven by (a) frequency of soil combination and mixing, or (b) frequency of vortex mixing in vortex controls. The community assembly processes are assigned by a null modeling approach: the influence of selection is determined by comparing the observed *β*-mean nearest taxon distance (*β*MNTDObs; the mean phylogenetic distance between each OTU in one tube and its closest relative in another tube within the same mixing set) to the *β*MNTDNull, a null distribution of *β*MNTD values amongst 1× controls. The influence of dispersal is determined by comparing the observed Bray-Curtis dissimilarity (BC-DisObs) between two tubes within the same mixing set to the BC-DisNull, a null distribution of BC-Dis values. Both bMNTDNull and BC-DisNull were calculated amongst the 32 1× controls when randomly assigned to groups of eight, iterated 999 times. Error bars represent ± 1 standard deviation.

The vortex controls, which were stand-alone tubes of soil that underwent vortex mixing but were never combined with other soil, were generally undominated by any particular community assembly processes (Fig. 6b), particularly at lower frequencies of vortex mixing. At 32× and 16×, almost 50% of the pairwise comparisons demonstrated dispersal limitation and/or variable selection; these community assembly processes characterize community pairs that are more dissimilar or have higher phylogenetic turnover than the null model does (created from 1× controls).

### Taxonomic composition shifted with increased soil mixing

In order to better understand community coalescence and to identify key taxa associated with community assembly processes in soil, we explored shifts in community composition related to the soil mixing treatments. The 1× controls had the highest phylum-level relative abundances of *Proteobacteria*, *Acidobacteria*, *Chloroflexi*, *Verrucomicrobia*, and *Actinobacteria*, which comprised over 80% of mean relative abundance and reflected the phylum-level composition of the initial soil communities (SI Fig. 5). After frequent soil mixing, over 80% of mean relative abundance were taxa from the phyla *Actinobacteria* and *Proteobacteria*, with one genus, *Nocardioides* (*Propionibacteriales*), comprising almost 30% of the mean relative abundance at 32×, and with one particular OTU emerging as the most abundant OTU in each of the 32× communities. Also within *Actinobacteria*, the second most abundant OTU at 32× was identified as belonging to the family *Micrococcacae*, with numerous 100% identity matches on NCBI including taxa from the genera *Arthrobacter*, *Paenarthrobacter*, and *Pseudarthrobacter* and comprising almost 10% of the mean relative abundance at 32×. The most abundant *Proteobacteria* at 32× were primarily comprised of *Rudaea* and *Rhodanobacter* (both *Gammaproteobacteria*), which together comprised over 15% of the mean relative abundance at 32×. The dominance of several OTUs at high soil mixing frequencies is apparent in the stark difference in cumulative mean relative abundance curves for the unmixed vs. frequently mixed treatments (SI Fig. 6), in which the four most abundant OTUs at 32× (named above), and the ten most abundant OTUs at 16× comprised over 50% of the cumulative mean relative abundance; this same proportion was comprised of almost 100 OTUs in the infrequently or unmixed treatments. Unlike their mixed soil counterparts, the vortex controls generally resembled the 1× controls in their mean relative abundances of phyla across vortex mixing frequencies (SI Fig. 5).

When assessing taxonomic differential abundance relative to 1× controls, we found 392 taxa with positive differential abundance (i.e. enriched in mixing treatments). There were 77, 202, 223, 171 and 180 enriched taxa in the 2×, 4×, 8×, 16×, and 32× treatments, respectively. To make our assessment more tractable, we focused on taxa with the biggest response (μ > 1.0), and only considered enriched taxa with mean relative abundances greater than 0.002 (0.2%) following enrichment. Fewer than 10% of the total enriched taxa met these criteria, leaving 5, 11, 18, 20, and 24 strongly enriched and common taxa in each treatment, respectively (SI Table 1, SI Fig. 7). These taxa were mostly from the phyla *Actinobacteria* and *Proteobacteria*. The most relatively enriched OTU (highest differential abundance estimate) at 32× relative to 1× was from the family *Nocardioidaceae (Actinobacteria)* and had no 100% ID matches in the NCBI nucleotide database. There were four additional OTUs from the genus *Nocardioides* that were considered relatively enriched, including the most abundant OTU found in every 32× tube, referenced above. In this case, it is possible that these similar OTUs are, in fact, different copies of the rRNA gene that exist within a singular organism’s genome, and this is an instance of splitting a single genome into separate OTUs (Schloss 2021). The second most relatively enriched OTU at 32× was from the genus *Lysobacter (Proteobacteria)*. Though some OTUs increased in relative abundance monotonically with increasing soil mixing frequency (e.g. OTUs from *Nocardioides* and *Oryzihumus*; SI Fig. 8), other taxa peaked in relative abundance at moderate frequencies of soil mixing (e.g. OTUs from *Rhodanobacter* and *Microscillaceae*; SI Fig. 9). One OTU from the genus *Nevskia* (*Xanthomonadaceae*) was relatively enriched at 2× and 4×, but relatively depleted at 32× (SI Fig. 9; this OTU had the same trend in the vortex controls).

There were 2152 total taxa with negative differential abundance (i.e. depleted) in the mixed treatments compared to 1×, which greatly exceeded the number of enriched taxa. By treatment, there were 66, 479, 1379, 1794, and 1993 depleted taxa for the 2×, 4×, 8×, 16×, and 32× treatments, respectively. Similarly to our approach for enriched taxa, we focused on the taxa with the strongest response (μ < −1.0), and only retained depleted taxa that were not extremely rare to begin with (mean relative abundances greater than 0.002 in the 1× treatment). This approach retained just 3, 17, and 40 strongly-depleted and common taxa in the 8×, 16×, and 32× treatments (SI Table 2, SI Fig. 10), and no taxa from the 2× and 4× treatments. These taxa were mostly from the phyla *Acidobacteria*, *Proteobacteria*, and *Verrucomicrobia*, and monotonically decreased in relative abundance with increased frequency of mixing (e.g. *Ktedonobacteraceae* and *Xanthobacteraceae*; SI Fig. 11), with some taxa maintaining relative abundances through the 2× and 4× mixing rates and becoming depleted with increased mixing frequency (e.g. *Acidothermus* and an OTU from the family *Streptomycetaceae*).

In the vortex controls, we found a similar number of enriched taxa with positive differential abundance, and some overlap with the OTUs found to be enriched in the mixed soil treatments (SI Fig. 12), but notably fewer depleted taxa (SI Fig. 13), with just four OTUs depleted across treatments, after filtering out the very rare or weakly responding taxa as described above. These four OTUs were also depleted in the mixed soil treatments.

### Predicted weighted mean 16S rRNA gene copy number increased with mixing

The weighted mean predicted 16S rRNA gene copy number was statistically different across mixing treatments (ANOVA, *p* < 0.001), and increased with mixing frequency from a mean value of 2.09 for 1× to a value of 2.51 for 32× (SI Fig. 14a). Each treatment, except 2×, was significantly higher than 1× (*p* < 0.05 for 4×; *p* < 0.001 for 8×, 16×, and 32×). Notably, the proportion of OTUs for which a predicted 16S rRNA gene copy number was available also increased with rate of mixing; about 30% of 1× OTUs as compared to 64% of 32× OTUs had matching genera in the rrnDB. *Nocardioides sp*., which comprises over 30% of relative abundance in 32×, largely accounts for this difference in OTU copy number availability by treatment (SI Table 3). Further, with a predicted mean copy number of 2.62, *Nocardioides sp*. heavily weighs on this analysis, given the high proportion of OTUs for which we don’t have a predicted gene copy number. Thus, we tested the degree of influence of *Nocardioides* on this analysis by simply removing its weighted mean predicted gene copy number from our calculations and found that the trends remained significant (ANOVA, *p* < 0.001), though to a slightly lesser extent at 8×, 16×, and 32× where *Nocardioides* is particularly relatively abundant (SI Fig. 14b and SI Table 3).

### Co-Occurrence Network Analysis

We created compositional networks to consider co-occurrences and co-exclusions of OTUs within each treatment. Co-occurrence network properties are reported in SI Table 4. Network figures, relative abundance of OTUs within each module by mixing set, and relative abundance of OTUs within each module across all mixing treatments are presented in SI Figs. 16–22. The initial network (SI Fig. 16), representing the communities captured by random sampling of freshly collected soil, has low modularity (0.24; SI Table 4) and is comprised of just 5 OTUs (nodes), or < 0.1% of the total taxa across initial samples, and < 0.05% of relative abundance. Modularity is also low at 1×, but we do see modules of co-occurring taxa (e.g. modules 1 and 4; SI Fig. 17a), that are also co-excluded with other taxa (primarily found in module 3), which indicates that these undisturbed tube communities did have some compositional patterns. Modularity generally increases with soil mixing, with a higher proportion of the community included in the networks of frequently mixed soil (SI Table 4). Mixing also gives rise to modules characteristic of a specific mixing sets. For instance, at 8× we see that over 40% of the relative abundance in mixing set 1 is found in taxa from module 4, and that this mixing set has lower relative abundances of taxa in the other modules, which are more representative of the other mixing sets (SI Fig. 20b). This is also reflected in the numerous co-exclusions between module 4 and modules 2 and 3 (SI Fig. 20a). Mixing sets at 32× also demonstrate module specificity; taxa from module 1 are clearly characteristic of mixing set 2, whereas taxa from module 3 are dominant in the other mixing sets.

## Discussion

The aim of this study was to determine the effects of coalescence on community composition and ecological community assembly processes in the patchy, disconnected, heterogeneous soil environment. Consistent with our hypothesis, we found that more frequently mixed soil harbored less rich bacterial communities and, as we predicted, soil in the more frequently mixed sets demonstrated evidence for homogenizing dispersal and homogeneous selection. The findings from this study impact our understanding of how physical disturbance affects soil communities and contribute to our growing understanding of the vast bacterial diversity observed in soil.

### Bacterial richness and community coalescence

Using soil subsamples that would generally be considered homogeneous and highly similar in community composition (see ‘Initial’ samples in Fig. 4), we demonstrated the effects of community coalescence across a range of soil mixing frequencies. Frequently mixed soil had 20% lower bacterial richness compared to the 1× control (Fig. 3), which may be attributable to competitive exclusion under changing resource availability, as may be suggested by co-excluded taxa (SI Figs. 20a, 21a, 22a), or attributable to selection for stress-resistant organisms when abiotic conditions shifted beyond organismal tolerance (Rillig *et al.* 2015; Castledine *et al.* 2020). These results closely mirror those of a meta-analysis (Rocca *et al.* 2019), which found alpha diversity in soil decreased by a mean of 20% across a variety of environmental disturbances that encompassed a range of stressors, with conditional effects. For example, heat shock in agricultural soils decreased richness, but warming of permafrost soils increased richness (Rocca *et al.* 2019). In our study, some combination of stress and competition could reasonably decrease richness, and being a closed system, we would not anticipate sources of increased richness (from speciation, for example) over the relatively short incubation interval.

The variable selection observed across the vortex controls (Fig. 6b) not only helps to explain their maintenance of richness (Fig. 3), but also supports the possibility that dissimilarity amongst these closed communities may have been driven by heterogeneous resource availability. For instance, an idiosyncratic fragment of organic matter in one tube could contribute to the rapid growth and selection for a particular community. As a vortex control, this tube would be an isolated community that registers as variable selection when compared to its analogous controls. Conversely, if the tube belonged to a mixing set, this community would be subsequently dispersed throughout the soil in all tubes of the mixing set, thus decreasing within-set dissimilarity (as seen in Fig. 5b) and likely contributing to both homogeneous selection and homogenizing dispersal (Fig. 6a). The vortex controls may be analogous to soil aggregates, which can host isolated communities under variable selection due to patchy resource availability (Wilpiszeski *et al.* 2019). While we would expect that some microbes may continue to remain isolated in protected soil pore spaces, or manage to persist due to priority effects during coalescence events (Castledine *et al.* 2020), our results suggest that, under frequent mixing conditions, the swift ascendency of a few taxa generally outweighs these effects, with a parallel outcome of decreased richness.

### Nocardioides thrive with soil mixing

Through increased connectedness, forced chance encounters, and potentially degraded refuges, frequently mixed soil becomes depleted in some taxa and relatively enriched in others. With soil mixing, new communities developed that were distinct and dissimilar from the 1× controls (Figs. 4 and 5), and initially rare OTUs become abundant after repeated community coalescence events (SI Fig. 8). These apparently mixing-loving, or at least mixing-tolerant organisms are likely generalists that translate available resources into fast growth rates; similar results were found in a coalescence experiment targeting life strategy in a lagoon, where a diverse bacterial community of oligotrophic specialists was overcome by copiotrophic generalists (Sara Beier, ESA 2021). Under coalescence, several OTUs of the genus *Nocardioides* (potentially representing the same organism – see note in the *Results* section) emerged as the most relatively enriched and abundant OTUs in frequently mixed treatments (SI Figs. 7 & 8). This is supported by previous work that found *Nocardioides* to be relatively enriched in both mixed soil treatments of a reciprocal coalescence study (A added to B or B added to A where *Nocardioides* originated in A, an industrial waste soil) (Wu *et al.* 2019), and also enriched in earthworm drilosphere soil compared to bulk soil in a no-tillage wheat experiment (i.e. the high disturbance hotspots in an otherwise undisturbed soil environment) (Schlatter *et al.* 2019). Another study found that emergent rare taxa comprised over half of the observed OTUs in the coalescent communities in mixed brackish waters, with many of these rare taxa becoming highly abundant (Rocca *et al.* 2020). The genus *Nocardioides* has also been associated with straw mineralization (Bernard *et al.* 2012), extracellular DNA degradation (Morrissey *et al.* 2015), rapid atrazine mineralization (Topp *et al.* 2000), and increased abundance after nanomaterial addition in maize (Zhang *et al.* 2020). *Nocardioides* has a predicted mean 16S rRNA gene copy number of 2.62 (rrnDB; Stoddard et al., 2015), which is higher than the community mean copy numbers predicted for either the initial (unincubated) community or the 1× control in this study, which were both just over 2, though not particularly high compared to copy numbers measured in targeted growth rate studies (Klappenbach, Dunbar and Schmidt 2000; Nemergut *et al.* 2016). The 16S rRNA gene copy number for *Nocardioides* contributes to the increase in mean predicted 16S rRNA gene copy number that is associated with increased mixing frequency (SI Fig. 14a), but this finding is not solely driven by the high abundance of *Nocardioides*: the gene copy number increased with mixing frequency even when *Nocardioides* OTUs were excluded from this calculation (SI Fig. 14b), indicating a mixing-driven proclivity for increased gene copy number beyond that of one organism or genus, and suggests that this trait has a selective advantage during frequent coalescence. Community coalescence is an often-overlooked form of disturbance (Mansour et al., 2018; e.g. Rocca et al., 2019), but it may help to describe the frequently-observed phenomena of emergence and enrichment of previously rare taxa under rapidly changing biotic and abiotic conditions (Allison and Martiny 2008).

### Mixed soil community assembly

Frequent soil mixing was associated with both deterministic and stochastic community assembly processes (homogeneous selection and homogenizing dispersal, respectively), and moderate mixing was associated with more stochastic assembly (Fig. 6a). This demonstrates that large populations (here, the OTUs that arise to high relative abundances with frequent mixing) tend to be governed by deterministic forces, and small or rare populations (here, rare taxa that persist in the less-frequently mixed sets) are more subject to stochasticity and drift (Hanson *et al.* 2012). However, specific community composition in a given tube was not driven by mixing frequency alone. Though we observed deterministic enrichment of *Nocardioides* OTUs and other strong responders with frequent mixing, there were also characteristic modules of community members in each frequently-mixed set (SI Figs. 20b, 21b, 22b), contributing to a high level of dissimilarity across mixing sets (Fig. 5c, see also Fig. 4) as opposed to the within-mixing set comparisons becoming more similar (Fig. 5b). This mixing set-specific response stands in contrast to the trend observed with moderate mixing, in which more stochastic processes observed at 2× and 4× produce a mixing set-agnostic response by which we see increasingly similar community composition whether comparisons are made within mixing sets (Fig. 5b) or outside of mixing sets (Fig. 5c), and where we also see more sparse co-occurrence networks (SI Figs. 18a, 19a). Furthermore, as mixing frequency increased, comparisons made across mixing sets remained similar to those at 1X (Fig. 5c). Together, this emphasizes that, while mixing soil likely selects for mixing-adapted taxa, the specific outcomes of community composition will differ, depending on small differences within starting communities.

Methodologically, our identification of community assembly processes represents a snapshot in time. For example, community sampling directly after soil mixing in a moderate treatment might be more likely to capture the effects of dispersal, as opposed to the community resulting from weeks of post-disturbance recovery. By the same token, it would also be interesting to sample the frequently-mixed community following a lengthy, undisturbed incubation. This ‘recovery’ period might favor more deterministic processes, as was observed in a chronosequence study that found a shift from stochastic forces characterizing early stages of succession, to deterministic homogeneous selection characterizing later stages of succession (Dini-Andreote *et al.* 2015).

### Vortex controls represent isolated communities

The vortex controls, which were individually mixed by vortex but never combined with other soil, are an informative and critical component of this research. Like the mixed treatment soil, soil within each vortex control tube is also undergoing coalescence, just on a smaller, within-tube scale, as the microhabitats and distinct communities within a tube are mixed to create new environments and assemblages of microbes. Community assembly processes in the vortex controls contrasted with our mixed soil treatments in that we detected both variable selection and dispersal limitation relative to the 1× control null model, and most comparisons were considered undominated by any particular community assembly process (Figs. 5b and 6). Critically, these results first highlight that the contrasting findings in the mixed sets are not simply due to the soil being physically agitated – rather, the findings discussed above are specifically the result of combining communities during mixing. However, we can also consider the results of the vortex controls on their own merit. If community assembly in mixed soil was driven solely by deterministic processes, homogenizing selection would have been the dominant process, producing isolated communities that are adapted to the physical effects of mixing and resulting in similar communities across the vortex controls within each mixing frequency (Evans, Martiny and Allison 2017). However, we instead observe some level of inherent stochasticity in these isolated soil communities (i.e. contained in separate tubes), that yields a collective of vortex control communities increasingly structured by dispersal limitation and variable selection as mixing frequency increased (Fig. 6b). Increased variable selection may be due to differences in tube-level resources (Wilpiszeski *et al.* 2019) that are accentuated through repeated within-tube coalescence. The dispersal-related processes are more surprising, though, since each vortex control tube, regardless of the number of times vortex mixed, was similarly isolated from all other tubes (e.g. the 2× vortex controls were just as isolated from each other as the 32× vortex controls). With 30–40% of comparisons demonstrating dispersal limitation at 16× and 32× (Fig. 6b), communities under more frequent within-tube coalescence via mixing presumably change faster and thus differentiate more over the 16-week incubation period, thus making dispersal limitation a more distinct community assembly process compared to moderate vortex mixing. This may be attributable to more frequent interactions amongst organisms that were not interacting in the 1× controls, or the effects of soil mixing on extracellular DNA, such as increased feeding or decreased organo-mineral associations (see discussion below in *Methodological considerations*). Another notable observation lies in the comparison between the 2× mixed soil treatment and the 2× vortex controls: these tubes only differed in their treatment and handling at one single mixing event, halfway through the incubation, during which the 2× mixed soil treatment was mixed within sets, and the 2× vortex controls underwent only within-tube vortex mixing. However, we see a sizeable difference in the outcome, with almost 30% of pairwise observations in the mixed soil treatment demonstrating homogeneous selection and/or homogenizing dispersal (Fig 6a), whereas the 2× vortex controls are largely undominated (Fig. 6b). This highlights how one soil mixing event was sufficient to flip undominated outcomes (and possibly some dispersal limitation and variable selection) into homogenizing processes. Further, this comparison suggests that even subtle or infrequent soil coalescence events, such as annual tillage or soil moved during construction or restoration, could substantially shift community composition and its driving processes.

### Disturbance disrupts mechanisms that maintain soil bacterial diversity

Generally speaking, soil is relatively undisturbed. That said, soils have fauna that burrow and consume soil. Root growth displaces soil, and subsequently root senescence creates pore space. Soil microbes themselves contribute to aggregate formation, organo-mineral associations, and other miniature soil “disturbances”. Similarly, cryoturbation and tillage can be significant across a landscape. These bigger disturbance events are fragmented, point disturbances, and occur perhaps only occasionally in any given location. In this experiment, we demonstrated that even infrequent soil coalescence can have an impact on community composition and community assembly processes, while frequent community coalescence events resulted in significant losses of bacterial richness and the introduction of deterministic selective processes. We expect the selective processes at work are likely biotic, as we see sharp increases in the relative abundances of likely copiotrophic bacteria such as *Nocardioides*, in the absence of typical environmental selection filters (e.g. pH, temperature, moisture). Our findings generally support the hypothesis that both soil heterogeneity and spatial dysconnectivity underpin the high diversity of the inhabitant microbial communities in soil (Fierer and Jackson 2006; Portell *et al.* 2018).

### Methodological considerations

While estimates of soil bacterial richness are notoriously hindered by constraints of sequencing depth given extremely high baseline richness, we tried to mitigate this by extracting DNA from the entire soil sample, sequencing deeply (mean 45 000 denoised reads per sample), and using appropriate algorithms to estimate richness (e.g. Willis and Bunge, 2014). Thus, we expect that our detection of decreased richness with mixing is not simply an artifact of a few taxa blooming in abundance.

The observed changes in relative abundances of OTUs indicate that compositional differences are due in large part to changes in the active bacterial community over the course of the experiment. However, we must also consider the effect of DNA from inactive (e.g. dead or dormant) bacteria, which is a constituent of total genomic DNA extracted from soils. Some research indicates a timescale of days to weeks for either rapid decomposition of much of the extracellular, or ‘relic’ DNA by the active community, or sorption to soil particles, such that this relic DNA evades extraction (Morrissey *et al.* 2015). Frequent mixing may inhibit formation of organo-mineral complexes between cellular components and soil particles, or may dislodge DNA from protected areas. As a result, soil disturbance likely suppresses detection of this less protected relic DNA due to heightened degradation, but disturbance may also enhance relic DNA detection if it is more chemically available for extraction due to weaker associations with soil. Overall, some work suggests that the relic DNA pool does not strongly influence richness estimates (Lennon *et al.* 2018), whereas another study demonstrates that relic DNA can inflate bacterial richness by an average of 14% across a variety of soils (Carini *et al.* 2016), though with a lesser effect in low pH soils, such as ours. Regardless, the strength of observed community changes in the frequently mixed treatments were large (Fig. 5), and not likely driven primarily by relic DNA.

In contrast to relic DNA, dormancy may play a larger role in affecting community composition, as it may account for half of OTUs found in soil, or 80% of cells (Lennon and Jones 2011). Had we been only interested in active taxa, we could have measured protein synthesis potential using rRNA, which is likely higher in active organisms, but this approach can be problematic in diverse communities and may still capture dormant organisms (Blazewicz *et al.* 2013). Though relative abundances of rRNA vary depending on activity, relative abundances of DNA are more static, and therefore a more appropriate target of measurement for community assembly processes. Overall, we expect that the inclusion of dead or dormant bacteria in our analyses is not particularly problematic because we were not assessing community function, but, instead, interpreting relative changes in the soil bacterial community composition. Methods to exclude extracellular DNA (Carini *et al.* 2016) or to isolate the active community (Romanowicz *et al.* 2016) would introduce other biases and interpretive challenges.

### Future directions

Soil functional assessment was beyond the scope of this work, so we are unable to expand upon the complicated and seemingly elusive relationships between microbial community diversity and function (Raynaud and Nunan 2014; Young and Bengough 2018). A future direction might be to assess microbial community function in soil undergoing natural coalescence events. We could predict that frequently mixed soil may exhibit decreased potential functional breadth due to decreased richness and the dominance of several OTUs that are adept at rapid growth following mixing. However, due to high functional redundancy in soil microbial communities (Louca *et al.* 2018), whether there would be meaningful impacts from such a reduction may be questionable. Further, rare taxa, which we found to be characteristic of frequent community coalescence (see also Allison and Martiny 2008) impart phylogenetic plasticity to the microbiome, which can enable functional resilience during periods of transition (Jousset *et al.* 2017; Jia, Dini-Andreote and Salles 2018), and therefore we might also predict that soil function is maintained despite a 20% decrease in richness with frequent mixing. To test function and community resilience, mixed and undisturbed soil communities could be coalesced to compare final community function under an array of environmental filters or carbon sources. Another direction could be to test spatially fragmented and isolated coalescence events, which could have high compositional and functional resilience in part due to proximity of the coalesced soil to undisturbed areas, or dilution of the outcomes. Finally, another extension of this work could be to study the effects of soil mixing on fungi, which play an important role in soil structure and function (Crawford *et al.* 2012); there are likely particular implications of disturbance by mixing for filamentous fungi that spread, forage, and connect habitats (Cairney 2005).

## Conclusions

Community coalescence in what may be considered a homogeneous soil demonstrates that the bacterial community can change considerably with mixing to reveal a strong dominance of several opportunistic OTUs, as bacterial richness otherwise declines. Homogeneous selection and homogenizing dispersal were the predominant community assembly processes in frequently mixed soil, whereas less disturbed soil was undominated by any particular community assembly processes and was therefore more subject to drift. On the other hand, variable selection and dispersal limitation were more apparent processes across isolated soil communities (here, the vortex controls). Our results generally suggest that soil heterogeneity, preserved in relatively unmixed soil, underpins the vast microbial diversity characteristic of soil.

## Supplementary Information

Supplementary Information is available online.

## Acknowledgements

We would like to acknowledge the contributions by Jamie Woolet and Dik Patterson towards making this work possible. This work was financially supported by the O.N. Allen Professorship (UW-Madison College of Agriculture & Life Sciences), awarded to Thea Whitman in 2016; the Louis and Elsa Thomsen Wisconsin Distinguished Graduate Fellowship (UW-Madison College of Agriculture & Life Sciences), awarded to Jaimie West in 2020; and a NSF EAGER grant (award # 2024230).

## References

Adler PB, Hillerislambers J, Levine JM. A niche for neutrality. Ecology Letters 2007;10:95–104.

Allison SD, Martiny JBH. Resistance, resilience, and redundancy in microbial communities. Proceedings of the National Academy of Sciences of the United States of America 2008;105:11512–9.

Anderson MJ. A new method for non-parametric multivariate analysis of variance. Austral Ecology 2001;26:32–46.

Anderson MJ. Distance-Based Tests for Homogeneity of Multivariate Dispersions. Biometrics 2006;62:245–53.

Armitage DW, Jones SE. How sample heterogeneity can obscure the signal of microbial interactions. The ISME journal 2019;13:2639–46.

Bailey VL, McCue LA, Fansler SJ et al. Micrometer-scale physical structure and microbial composition of soil macroaggregates. Soil Biology and Biochemistry 2013;65:60–8.

Bell G. Neutral Macroecology. Science 2001;293:2413–8.

Benjamini Y, Hochberg Y. Controlling the False Discovery Rate: A Practical and Powerful Approach to Multiple Testing. J Royal Statistical Soc Ser B Methodol 1995;57:289–300.

Bernard L, Chapuis-Lardy L, Razafimbelo T et al. Endogeic earthworms shape bacterial functional communities and affect organic matter mineralization in a tropical soil. Isme J 2012;6:213–22.

Blazewicz SJ, Barnard RL, Daly RA et al. Evaluating rRNA as an indicator of microbial activity in environmental communities: limitations and uses. Isme J 2013;7:2061–8.

Bolyen E, Rideout JR, Dillon MR et al. Reproducible, interactive, scalable and extensible microbiome data science using QIIME 2. Nature biotechnology 2019;37:852–7.

Bouyoucos GJ. Hydrometer Method Improved for Making Particle Size Analyses of Soils1. Agron J 1962;54:464–5.

Bray JR, Curtis JT. An Ordination of the Upland Forest Communities of Southern Wisconsin. Ecological Monographs 1957;27:325–49.

Bray RH, Kurtz LT. Determination of Total, Organic, and Available Forms of Phosphorus in Soils. Soil Sci 1945;59:39–46.

Cairney JWG. Basidiomycete mycelia in forest soils: dimensions, dynamics and roles in nutrient distribution. Mycol Res 2005;109:7–20.

Calderón K, Spor A, Breuil M-C et al. Effectiveness of ecological rescue for altered soil microbial communities and functions. Isme J 2017;11:272–83.

Callahan BJ, McMurdie PJ, Rosen MJ et al. DADA2: High-resolution sample inference from Illumina amplicon data. Nature Methods 2016;13:581–3.

Carini P, Marsden PJ, Leff JW et al. Relic DNA is abundant in soil and obscures estimates of soil microbial diversity. Nature Microbiology 2016:1–6.

Carson JK, Gonzalez-Quiñones V, Murphy DV et al. Low pore connectivity increases bacterial diversity in soil. Applied and environmental microbiology 2010;76:3936–42.

Castledine M, Sierocinski P, Padfield D et al. Community coalescence: an eco-evolutionary perspective. Philosophical Transactions of the Royal Society B: Biological Sciences 2020;375:20190252–10.

Chase JM, Leibold MA. Ecological Niches: Linking Classical and Contemporary Approaches. University of Chicago Press, 2003.

Chase JM, Leibold MA. Revising the niche concept: definitions and mechanistic models. University of Chicago Press, 2014, 19–50.

Connor N, Barberán A, Clauset A. Using null models to infer microbial co-occurrence networks. Plos One 2017;12:e0176751.

Crawford JW, Deacon L, Grinev D et al. Microbial diversity affects self-organization of the soil– microbe system with consequences for function. J Roy Soc Interface 2012;9:1302–10.

Csardi G, Nepusz T. The igraph software package for complex network research. InterJournal 2006;Complex Systems:1695.

Delgado-Baquerizo M, Oliverio AM, Brewer TE et al. A global atlas of the dominant bacteria found in soil. Science 2018;359:320–5.

Dini-Andreote F, Stegen JC, Elsas JD van et al. Disentangling mechanisms that mediate the balance between stochastic and deterministic processes in microbial succession. Proceedings of the National Academy of Sciences of the United States of America 2015;112:E1326–32.

Doane TA, Horwath WR. Spectrophotometric Determination of Nitrate with a Single Reagent. Analytical Letters 2003;36:2713–22.

Dunnett CW. A Multiple Comparison Procedure for Comparing Several Treatments with a Control. J Am Stat Assoc 1955;50:1096–121.

Evans S, Martiny JBH, Allison SD. Effects of dispersal and selection on stochastic assembly in microbial communities. The ISME Journal 2017;11:176–85.

Fierer N. Embracing the unknown: disentangling the complexities of the soil microbiome. Nature Publishing Group 2017:1–12.

Fierer N, Jackson RB. The diversity and biogeography of soil bacterial communities. Proceedings of the National Academy of Sciences 2006;103:626–31.

Fine PVA, Kembel SW. Phylogenetic community structure and phylogenetic turnover across space and edaphic gradients in western Amazonian tree communities. Ecography 2011;34:552–65.

Fodelianakis S, Washburne AD, Bourquin M et al. Microdiversity characterizes prevalent phylogenetic clades in the glacier-fed stream microbiome. Isme J 2021:1–10.

Gewin V. Beyond Neutrality—Ecology Finds Its Niche. PLoS biology 2006;4:e278.

Gittel A, Bárta J, Kohoutová I et al. Distinct microbial communities associated with buried soils in the Siberian tundra. Isme J 2014;8:841–53.

Grinnell J. The Niche-Relationships of the California Thrasher. Auk 1917;34:427–33.

Guillou CL, Prévost-Bouré NC, Karimi B et al. Tillage intensity and pasture in rotation effectively shape soil microbial communities at a landscape scale. Microbiologyopen 2019;8:e00676.

Hanson CA, Fuhrman JA, Horner-Devine MC et al. Beyond biogeographic patterns: processes shaping the microbial landscape. Nature Reviews Microbiology 2012;10:497–506.

Hubbell SP. The Unified Neutral Theory of Biodiversity and Biogeography (MPB-32). Princeton University Press, 2001.

Huber P, Metz S, Unrein F et al. Environmental heterogeneity determines the ecological processes that govern bacterial metacommunity assembly in a floodplain river system. Isme J 2020;14:2951–66.

Hutchinson GE. Concluding remarks. Cold Spring Harbor Symposia. Quantitative Biology 1957;22:415–27.

Jacquiod S, Puga-Freitas R, Spor A et al. A core microbiota of the plant-earthworm interaction conserved across soils. Soil Biology Biochem 2020;144:107754.

Jia X, Dini-Andreote F, Salles JF. Community Assembly Processes of the Microbial Rare Biosphere. Trends in microbiology 2018;26:738–47.

Jousset A, Bienhold C, Chatzinotas A et al. Where less may be more: how the rare biosphere pulls ecosystems strings. The ISME Journal 2017;11:853–62.

Kembel SW, Cowan PD, Helmus MR et al. Picante: R tools for integrating phylogenies and ecology. Bioinformatics 2010;26:1463–4.

Klappenbach JA, Dunbar JM, Schmidt TM. rRNA operon copy number reflects ecological strategies of bacteria. Appl Environ Microbiol 2000;66:1328–33.

König S, Köhnke MC, Firle A-L et al. Disturbance Size Can Be Compensated for by Spatial Fragmentation in Soil Microbial Ecosystems. Frontiers Ecol Evol 2019;7:290.

Kuzyakov Y, Blagodatskaya E. Microbial hotspots and hot moments in soil: Concept & review. Soil Biology and Biochemistry 2015;83:184–99.

Lennon JT, Jones SE. Microbial seed banks: the ecological and evolutionary implications of dormancy. Nature Publishing Group 2011;9:119–30.

Lennon JT, Muscarella ME, Placella SA et al. How, When, and Where Relic DNA Affects Microbial Diversity. Mbio 2018;9:e00637–18.

Louca S, Polz MF, Mazel F et al. Function and functional redundancy in microbial systems. Nature Ecology & Evolution 2018;2:936–43.

Lynch MDJ, Neufeld JD. Ecology and exploration of the rare biosphere. Nature Reviews Microbiology 2015;13:217–29.

Mansour I, Heppell CM, Ryo M et al. Application of the microbial community coalescence concept to riverine networks. Biol Rev 2018;93:1832–45.

Martin BD, Witten D, Willis AD. Corncob: Count Regression for Correlated Observations with the Beta-Binomial., 2021.

McMurdie PJ, Holmes S. phyloseq: An R Package for Reproducible Interactive Analysis and Graphics of Microbiome Census Data. Watson M (ed.). PLOS ONE 2013;8:e61217.

Morrissey EM, McHugh TA, Preteska L et al. Dynamics of extracellular DNA decomposition and bacterial community composition in soil. Soil Biology Biochem 2015;86:42–9.

Nemergut DR, Knelman JE, Ferrenberg S et al. Decreases in average bacterial community rRNA operon copy number during succession. Isme J 2016;10:1147–56.

Ning D, Yuan M, Wu L et al. A quantitative framework reveals ecological drivers of grassland microbial community assembly in response to warming. Nat Commun 2020;11:4717.

O’Brien SL, Gibbons SM, Owens SM et al. Spatial scale drives patterns in soil bacterial diversity. Environmental Microbiology 2016;18:2039–51.

Oksanen J, Blanchet FG, Friendly M et al. Vegan: Community Ecology Package., 2019.

Or D, Smets BF, Wraith JM et al. Physical constraints affecting bacterial habitats and activity in unsaturated porous media – a review. Advances in Water Resources 2007;30:1505–27.

Pasolli E, Filippis FD, Mauriello IE et al. Large-scale genome-wide analysis links lactic acid bacteria from food with the gut microbiome. Nat Commun 2020;11:2610.

Philippot L, Griffiths BS, Langenheder S. Microbial Community Resilience across Ecosystems and Multiple Disturbances. Microbiol Mol Biol R 2021;85, DOI: 10.1128/mmbr.00026-20.

Portell X, Pot V, Garnier P et al. Microscale Heterogeneity of the Spatial Distribution of Organic Matter Can Promote Bacterial Biodiversity in Soils: Insights From Computer Simulations. Frontiers in Microbiology 2018;9:1583.

Quast C, Pruesse E, Yilmaz P et al. The SILVA ribosomal RNA gene database project: improved data processing and web-based tools. Nucleic Acids Research 2013;41:D590–6.

Raynaud X, Nunan N. Spatial Ecology of Bacteria at the Microscale in Soil. Pappalardo F (ed.). PLOS ONE 2014;9:e87217.

R-Core-Team. R: A Language and Environment for Statistical Computing., 2018.

Richards LA. Diagnosis and Improvement of Saline and Alkali Soils. Soil Sci 1954;78:154.

Rillig MC, Antonovics J, Caruso T et al. Interchange of entire communities: microbial community coalescence. Trends in Ecology & Evolution 2015;30:470–6.

Rillig MC, Lehmann A, Aguilar-Trigueros CA et al. Soil microbes and community coalescence. Pedobiologia 2016;59:37–40.

Rillig MC, Muller LA, Lehmann A. Soil aggregates as massively concurrent evolutionary incubators. The ISME Journal 2017;11:1943–8.

Rocca JD, Simonin M, Bernhardt ES et al. Rare microbial taxa emerge when communities collide: freshwater and marine microbiome responses to experimental mixing. Ecology 2020;101:32–14.

Rocca JD, Simonin M, Blaszczak JR et al. The Microbiome Stress Project: Toward a Global Meta-Analysis of Environmental Stressors and Their Effects on Microbial Communities. Front Microbiol 2019;9:3272.

Romanowicz KJ, Freedman ZB, Upchurch RA et al. Active microorganisms in forest soils differ from the total community yet are shaped by the same environmental factors: the influence of pH and soil moisture. Fems Microbiol Ecol 2016;92:fiw149.

Schlatter DC, Reardon CL, Johnson-Maynard J et al. Mining the Drilosphere: Bacterial Communities and Denitrifier Abundance in a No-Till Wheat Cropping System. Front Microbiol 2019;10:1339.

Schloss PD. Amplicon Sequence Variants Artificially Split Bacterial Genomes into Separate Clusters. Msphere 2021;6:e00191–21.

Schulte EE, Hopkins BG. Estimation of Soil Organic Matter by Weight Loss-on-Ignition. In: Magdoff FR, Tabatabai MA, Hanlon EA (eds.). Soil Organic Matter: Analysis and Interpretation. Madison, WI: Soil Science Society of America, 1996, 21–31.

Sierocinski P, Milferstedt K, Bayer F et al. A Single Community Dominates Structure and Function of a Mixture of Multiple Methanogenic Communities. Curr Biol 2017;27:3390–3395.e4.

Stegen JC, Lin X, Fredrickson JK et al. Quantifying community assembly processes and identifying features that impose them. The ISME journal 2013;7:2069–79.

Stegen JC, Lin X, Fredrickson JK et al. Estimating and mapping ecological processes influencing microbial community assembly. Frontiers in Microbiology 2015;6, DOI: 10.3389/fmicb.2015.00370.

Stegen JC, Lin X, Konopka AE et al. Stochastic and deterministic assembly processes in subsurface microbial communities. The ISME Journal 2012;6:1653–64.

Stoddard SF, Smith BJ, Hein R et al. rrnDB: improved tools for interpreting rRNA gene abundance in bacteria and archaea and a new foundation for future development. Nucleic Acids Res 2015;43:D593–8.

Tecon R, Or D. Biophysical processes supporting the diversity of microbial life in soil. FEMS Microbiology Reviews 2017;41:599–623.

Thomas GW. Exchangeable Cations. Methods of Soil Analysis, Part 2 Chemical and Microbiological Properties. 1982, 159–65.

Topp E, Mulbry WM, Zhu H et al. Characterization of S-Triazine Herbicide Metabolism by a Nocardioides sp. Isolated from Agricultural Soils. Appl Environ Microb 2000;66:3134–41.

Treves DS, Xia B, Zhou J et al. A Two-Species Test of the Hypothesis That Spatial Isolation Influences Microbial Diversity in Soil. Microbial Ecology 2003;45:20–8.

Tripathi BM, Stegen JC, Kim M et al. Soil pH mediates the balance between stochastic and deterministic assembly of bacteria. Isme J 2018;12:1072–83.

Vellend M. Conceptual Synthesis in Community Ecology. The Quarterly Review of Biology 2010;85:183–206.

Vos M, Wolf AB, Jennings SJ et al. Micro-scale determinants of bacterial diversity in soil. FEMS Microbiology Reviews 2013;37:936–54.

Walters W, Hyde ER, Berg-Lyons D et al. Improved Bacterial 16S rRNA Gene (V4 and V4-5) and Fungal Internal Transcribed Spacer Marker Gene Primers for Microbial Community Surveys. Bik H (ed.). mSystems 2016;1:e00009–15.

Whitman TL, Whitman E, Woolet J et al. Soil bacterial and fungal response to wildfires in the Canadian boreal forest across a burn severity gradient. Soil Biology and Biochemistry 2019:107571–59.

Wickham H. Ggplot2: Elegant Graphics for Data Analysis., 2016. Willis AD, Bunge J. Package Breakaway., 2014.

Wilpiszeski RL, Aufrecht JA, Retterer ST et al. Soil Aggregate Microbial Communities: Towards Understanding Microbiome Interactions at Biologically Relevant Scales. Müller V (ed.). Applied and environmental microbiology 2019;85:689.

Wu X, Li J, Ji M et al. Non-synchronous Structural and Functional Dynamics During the Coalescence of Two Distinct Soil Bacterial Communities. Front Microbiol 2019;10:1125.

Yilmaz P, Parfrey LW, Yarza P et al. The SILVA and “All-species Living Tree Project (LTP)” taxonomic frameworks. Nucleic Acids Research 2013;42:D643–8.

Young IM, Bengough AG. The search for the meaning of life in soil: an opinion. European Journal of Soil Science 2018;69:31–8.

Zhang W, Jia X, Chen S et al. Response of soil microbial communities to engineered nanomaterials in presence of maize (Zea mays L.) plants. Environ Pollut 2020;267:115608.

Zhou J, Ning D. Stochastic Community Assembly: Does It Matter in Microbial Ecology? Microbiology and molecular biology reviews : MMBR 2017;81:1–32.

