## Supplementary_Information for "Disturbance by soil mixing decreases microbial richness and supports homogenizing community assembly processes"

#### DNA extraction, PCR amplification, and 16S rRNA library prep

The DNeasy PowerLyzer PowerSoil Kit (Catalog No. 12855, Qiagen, Germantown, MD) was used following manufacturer instructions with several modifications. First, quantity of soil in each incubation tube was at most 55 mg, which is less than the standard 250 mg. Thus, care was taken to transfer all soil and residue from the incubation tube to the PowerBead Tube, including washing the tube three times, each with 250 uL of the PowerBead Solution and vortex, to retrieve as much soil residue and DNA as possible. The incubation blanks, which consisted of one empty tube associated with each mixing set in the 1× and 32× treatments, with air holes drilled as for incubation tubes. were included to track potential general contamination over the course of the experiment, and were also washed in the same manner to extract any contaminating DNA. Further, every round of extraction contained at least one extraction blank, and the final extraction for each kit was also a blank. All extracted DNA was stored at or below −20 °C throughout stages of sequencing.

16S rRNA genes of extracted DNA were amplified in triplicate using PCR. Variable region V4 of the 16S rRNA gene was targeted using forward primer 515f and reverse primer 806r (Walters et al., 2016). Primers also contained barcodes and Illumina sequencing adapters (Kozich et al., 2013). The following reagents comprised each 25 µL PCR reaction: 12.5 µL Q5 Hot Start High-Fidelity 2X Master mix (Catalog No. M0494, New England BioLabs, Ipswich, MA), 1.25 µL 515f forward primer (10 mM), 1.25 µL 806r reverse primer (10 mM), 1 µL DNA extract, 1.25 µL Bovine Serum Albumin (Catalog No. 40220056, bioWORLD, Dublin, OH), and 7.75 µL PCR-grade water. The plate was sealed and briefly centrifuged prior to 30 PCR cycles on an Eppendorf Mastercycler nexus gradient thermal cycler (Hamburg, Germany) using the following parameters: 98 °C for 2 min + 30 × (98 °C for 10 seconds + 58 °C for 15 seconds + 72 °C for 10 seconds) + 72 °C for 2 min and 4 °C hold.

Successful amplification was verified via gel electrophoresis using 1% TAE agarose gel and Invitrogen SYBR Safe DNA Gel Stain (Catalog No. S33102, ThermoFisher Scientific, Carlsbad, CA). Each well was loaded with 5 µL PCR product mixed with 1 µL Purple 6X Gel Loading Dye (Catalog No. B7025S, New England BioLabs, Ipswich, MA). Successful amplification of target base pair length was confirmed using a 2-Log DNA Ladder (0.1–10.0 kilobases; Catalog No. N3200, New England BioLabs, Ipswich, MA) in each gel. Gel ran for ninety minutes at 115V. A photo was taken when the gel was finished running and wells were checked visually for amplification.

The SequalPrep Normalization Plate Kit (Catalog No. A1051001, ThermoFisher Scientific, Carlsbad, CA) was used to normalize amplicon yields across samples using a limited binding capacity solid phase. Normalization was performed following kit instructions using 25 µL of the pooled triplicate PCR product, and yielded 20 µL of eluted DNA per sample, which was pooled following elution. The combined DNA library was concentrated using a SpeedVac Vacuum Concentrator System (ThermoFisher Scientific, Waltham, MA) prior to further DNA purification using Wizard SV Gel and PCR Clean-Up System (Catalog No. A9281, Promega Corporation, Madison, WI). Kit instructions were followed except the final nuclease-free water application was divided into 30 µL and 20 µL increments with the incubation and centrifuge steps after both additions.

#### SI Tables

**SI Table 1**. Taxa with positive differential abundance (with mean relative abundance > 0.002 (0.2%) and coefficient of differential abundance μ > 1.0). *See accompanying file SI_Table_1.csv.*

**SI Table 2**. Taxa with negative differential abundance (with mean relative abundance > 0.002 (0.2%) and coefficient of differential abundance μ < -1.0). *See accompanying file SI_Table_2.csv.*

**SI Table 3**. Mean weighted gene copy number and proportion of OTUs present in rrnDB (Stoddard et al., 2015), by soil mixing treatment.

| | | All OTUs | | WITHOUT *Nocardioides* OTUs | |
|---|---|---|---|---|---|
| **Frequency of mixing** | **Vortex Control** | **Proportion of OTUs with copy number in rrnDB** | **mean weighted copy number** | **Proportion of OTUs with copy number in rrnDB** | **mean weighted copy number** |
| Initial | *na* | 0.305 | 2.07 | 0.305 | 2.07 |
| 1× | *na* | 0.304 | 2.09 | 0.304 | 2.09 |
| 2× | N | 0.313 | 2.06 | 0.309 | 2.06 |
| 4× | N | 0.332 | 2.13 | 0.314 | 2.11 |
| 8× | N | 0.444 | 2.27 | 0.325 | 2.16 |
| 16× | N | 0.493 | 2.35 | 0.276 | 2.17 |
| 32× | N | 0.639 | 2.51 | 0.351 | 2.43 |
| 2× | Y | 0.312 | 2.02 | 0.309 | 2.02 |
| 4× | Y | 0.34 | 2.04 | 0.329 | 2.03 |
| 8× | Y | 0.344 | 2.07 | 0.325 | 2.05 |
| 16× | Y | 0.388 | 2.15 | 0.335 | 2.09 |
| 32× | Y | 0.399 | 2.25 | 0.341 | 2.2 |

**SI Table 4**. Co-occurrence network properties for each mixing treatment. Isolated (or unconnected) network components were removed from the network prior to calculations, but the 2× and 4× networks contained large secondary components (2°), for which network properties were also calculated. Nodes = number of OTUs in the network, edges = number significant pairwise correlations, modules = number of modules in a network, modularity = extent to which nodes cluster into modules with edges occurring at a higher rate than expected at random, degree = number of connections (i.e. edges) for each OTU, path length is a measure of compactness, clustering coefficient is the network's frequency of three-way interactions, diameter is maximal-length shortest path among pairs of nodes. Also included in this table is a sense for extent of OTU membership (number and proportion) in each network, as a function of total community, and following filtering to relative abundance >0.005 across the dataset.

| **Frequency of mixing** | **nodes** | **edges** | **modules** | **modularity** | **degree (ave)** | **path length (ave)** | **clustering coeff** | **clustering coefficient (ave)** | **closeness** | | **edge density** | **dia.** | **total community** | | **cutoff 0.005** | | **# of components removed (# nodes)** |
|---|---|---|---|---|---|---|---|---|---|---|---|---|---|---|---|---|---|
| | | | | | | | | | **ave** | **sd** | | | **n.taxa** | **%** | **n.taxa** | **%** | |
| Initial | 5 | 6 | 2 | 0.24 | 2.40 | 1.40 | 0.55 | 0.75 | 0.185 | 0.042 | 0.60 | 2 | 5645 | 0.09 | 2382 | 0.21 | 5 (2, 2, 2, 2, 2) |
| 1× | 35 | 143 | 4 | 0.19 | 8.17 | 1.97 | 0.56 | 0.77 | 0.016 | 0.003 | 0.24 | 4 | 5200 | 0.67 | 2419 | 1.45 | 3 (3, 2, 2) |
| 2× | 14 | 22 | 3 | 0.38 | 3.14 | 2.44 | 0.48 | 0.71 | 0.033 | 0.007 | 0.24 | 5 | 5199 | 0.27 | 2436 | 0.57 | 5 (2, 2, 2, 2, 2) |
| 2× (2°) | 8 | 7 | 2 | 0.36 | 1.75 | 2.54 | 0.00 | 0.00 | 0.059 | 0.013 | 0.25 | 5 | 5199 | 0.15 | 2436 | 0.33 | |
| 4× | 17 | 21 | 5 | 0.42 | 2.47 | 3.00 | 0.22 | 0.22 | 0.022 | 0.005 | 0.15 | 6 | 4955 | 0.34 | 2434 | 0.70 | 4 (8, 5, 4, 2) |
| 4× (2°) | 13 | 17 | 4 | 0.46 | 2.62 | 2.59 | 0.28 | 0.24 | 0.034 | 0.008 | 0.22 | 6 | 4955 | 0.26 | 2434 | 0.53 | |
| 8× | 117 | 487 | 6 | 0.54 | 8.32 | 3.38 | 0.44 | 0.57 | 0.003 | 0.001 | 0.07 | 10 | 4850 | 2.41 | 2443 | 4.79 | 1 (2) |
| 16× | 106 | 1336 | 3 | 0.29 | 25.21 | 1.96 | 0.62 | 0.73 | 0.005 | 0.001 | 0.24 | 4 | 4540 | 2.33 | 2413 | 4.39 | 1 (4) |
| 32× | 70 | 344 | 3 | 0.51 | 9.83 | 2.64 | 0.53 | 0.68 | 0.006 | 0.001 | 0.14 | 7 | 4727 | 1.48 | 2404 | 2.91 | none |

### SI Figures

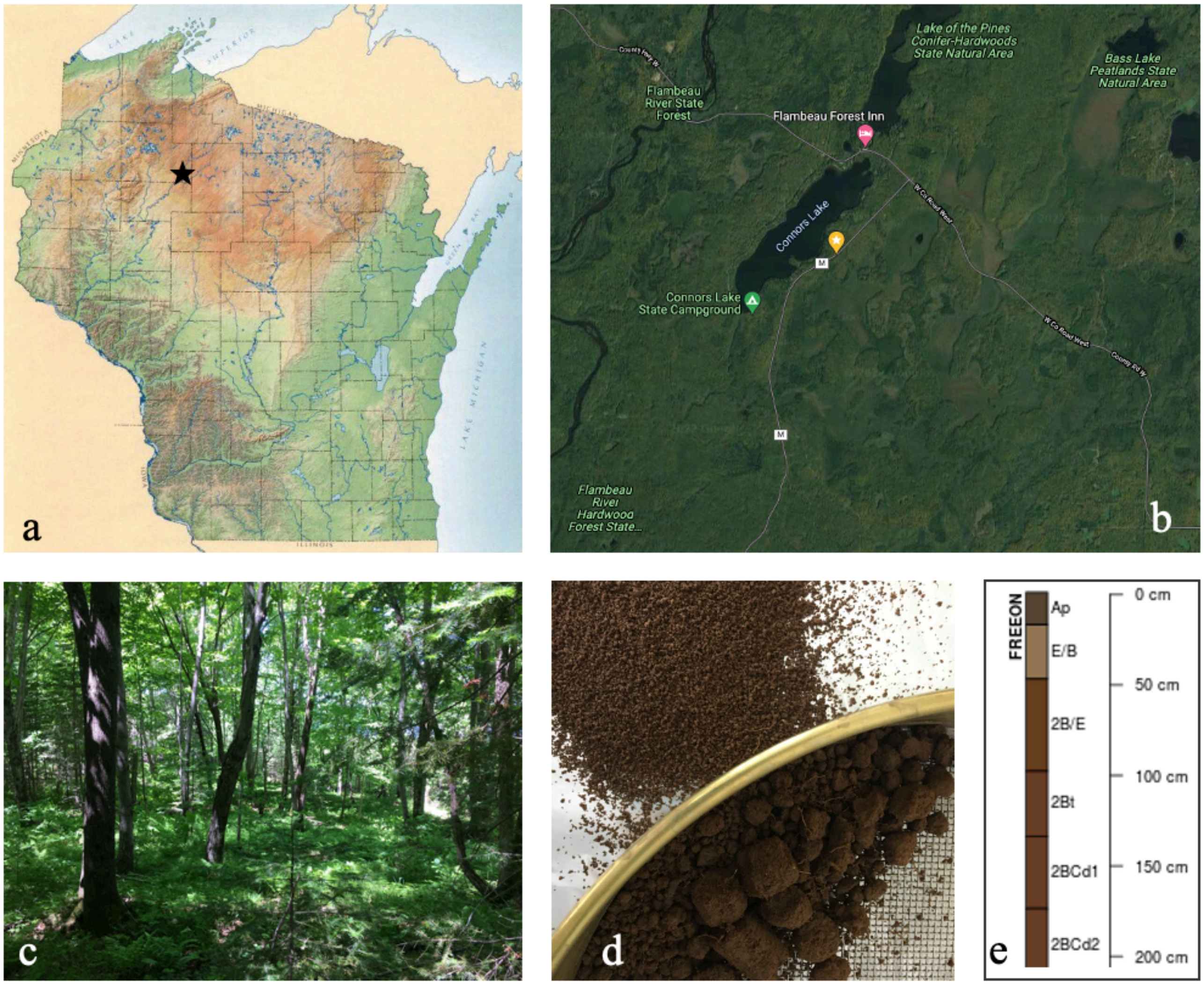

**SI Figure 1**. Overview of soil collection. **a & b**) Soil was collected on 30 August, 2018 near Connor's Lake in Sawyer County, Northern WI, U.S. (45°44'55.6"N, 90°43'47.1"W, 430 m asl). **c**) Vegetation type was northern mesic forest, early-to-mid seral, dominated by *Acer rubrum* L. (approx. 80%), *Acer saccharum* Marsh. (approx. 10%), *Betula alleghaniensis* Britt. (approx. 5%), and *Tilia americana* L. (<5%). **d**) Two soil cores (1.8 cm dia) were collected from each of 6 locations randomly chosen along a 50 m transect. From each of these 12 soil cores, we retained a portion of the A horizon, 15–20 cm of depth, and sieved this soil to 2 mm. **e**) Soil was Freeon silt loam soil, a very deep, moderately well drained, coarse-loamy, mixed, superactive, frigid Oxyaquic Glossudalf.

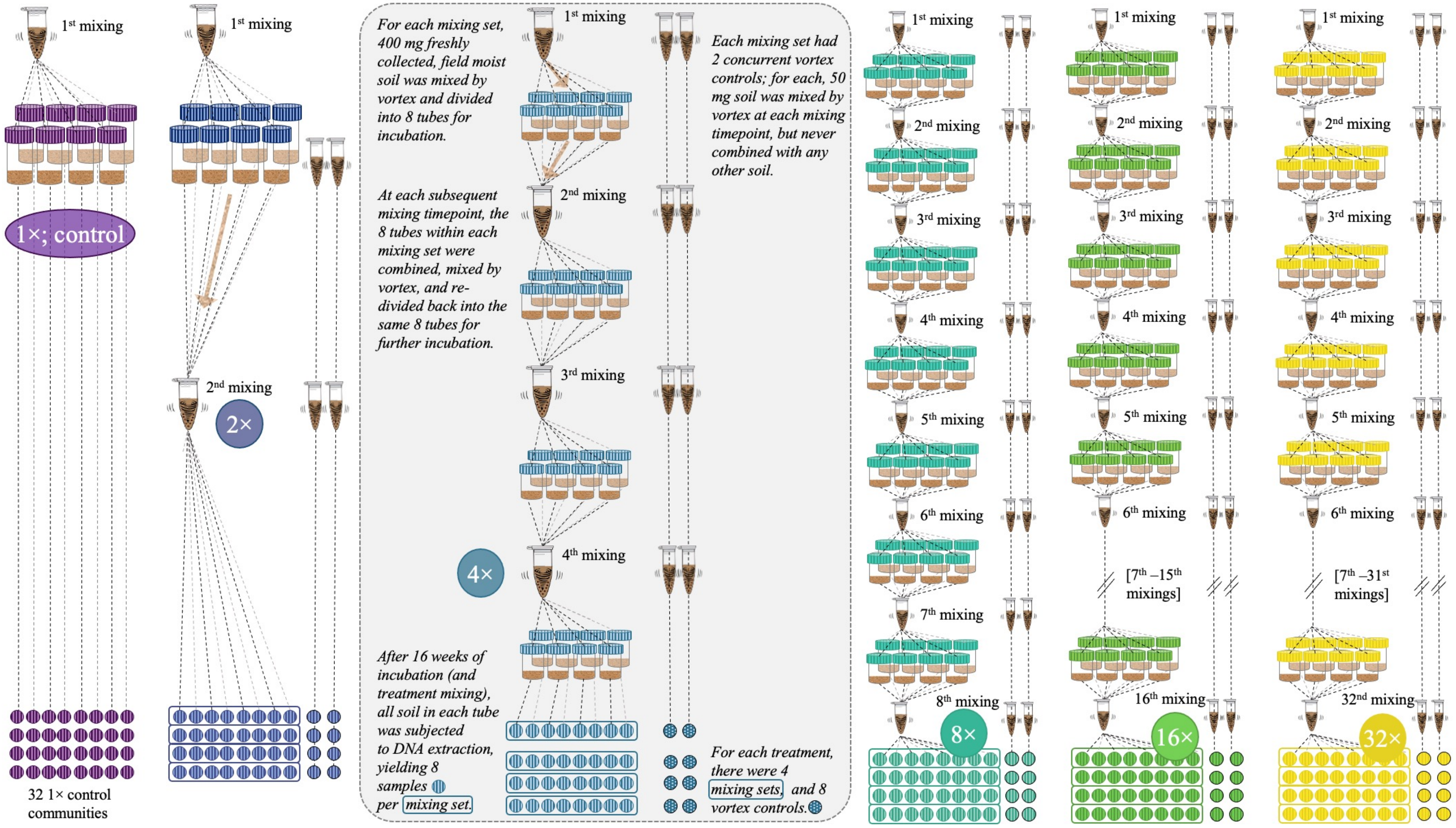

**SI Figure 2**. Experimental setup depicting one mixing set for each of the mixing treatments (2×—32×) plus 1× control. For each mixing set, 400 mg freshly collected soil was mixed by vortex and separated into 8 tubes, approximately 50 mg each. The tubes were incubated for regular intervals, between which the soil in all 8 tubes was combined, mixed by vortex, and re-divided back into the 8 tubes (note: the incubation interval varied in length depending on the mixing frequency, and 1× control tubes were not subject to further mixing). Concurrent vortex controls were stand-alone tubes of soil that were mixed by vortex at each mixing timepoint but never combined with other soil. At the conclusion of the 16-week incubation, DNA was extracted from soil in each incubation tube for 16S rRNA gene V4 region amplicon sequencing. Each treatment was comprised of 4 mixing sets (8 tubes in each) with 8 accompanying vortex controls.

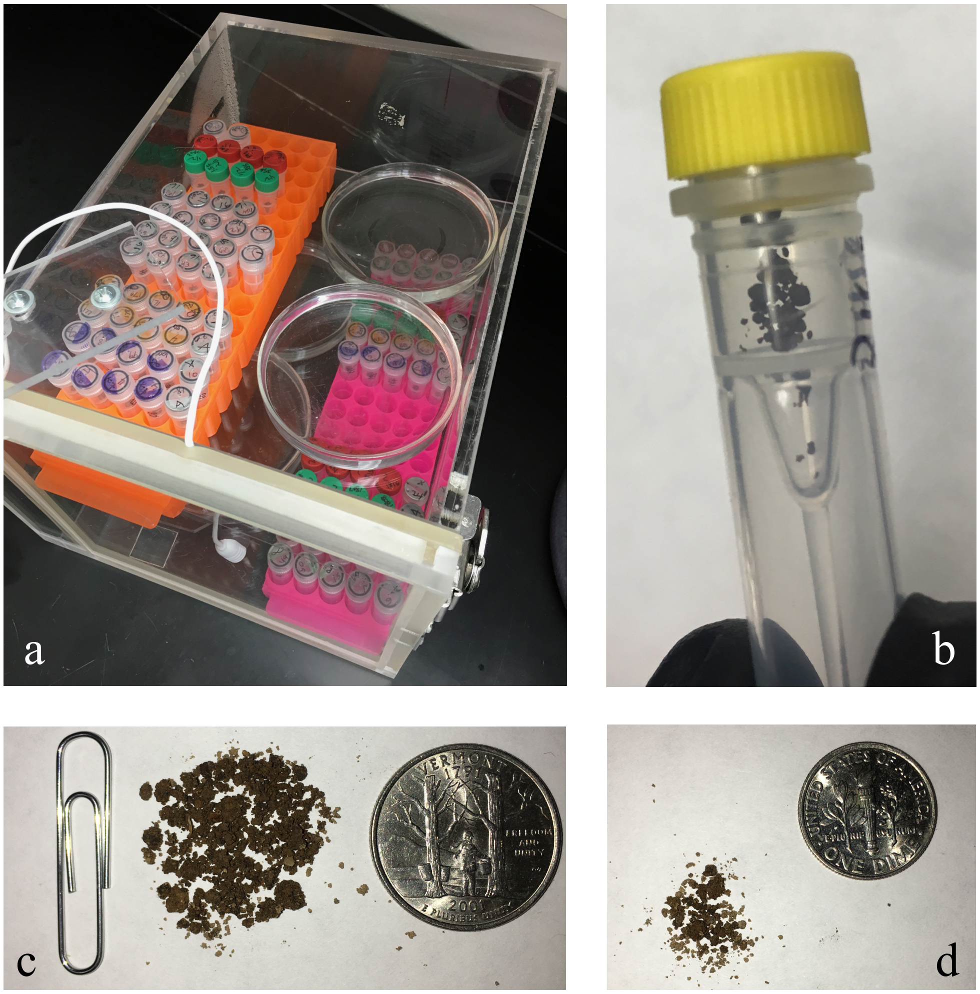

**SI Figure 3**. Laboratory incubation. **a**) Tubes of soil incubating in a sealed incubation box with open petri dishes of water to maintain relative humidity > 95%. The incubation boxes were covered for darkness. **b**) 50 mg soil in a 50 µL freestanding microcentrifuge tube (laying horizontal). **c**) 400 mg of soil, equivalent to one “mixing set”, visualized relative to a standard size paperclip and a U.S. quarter. **d**) 50 mg soil, equivalent to one tube, visualized next to a U.S. dime. c) and d) are shown at the same scale.

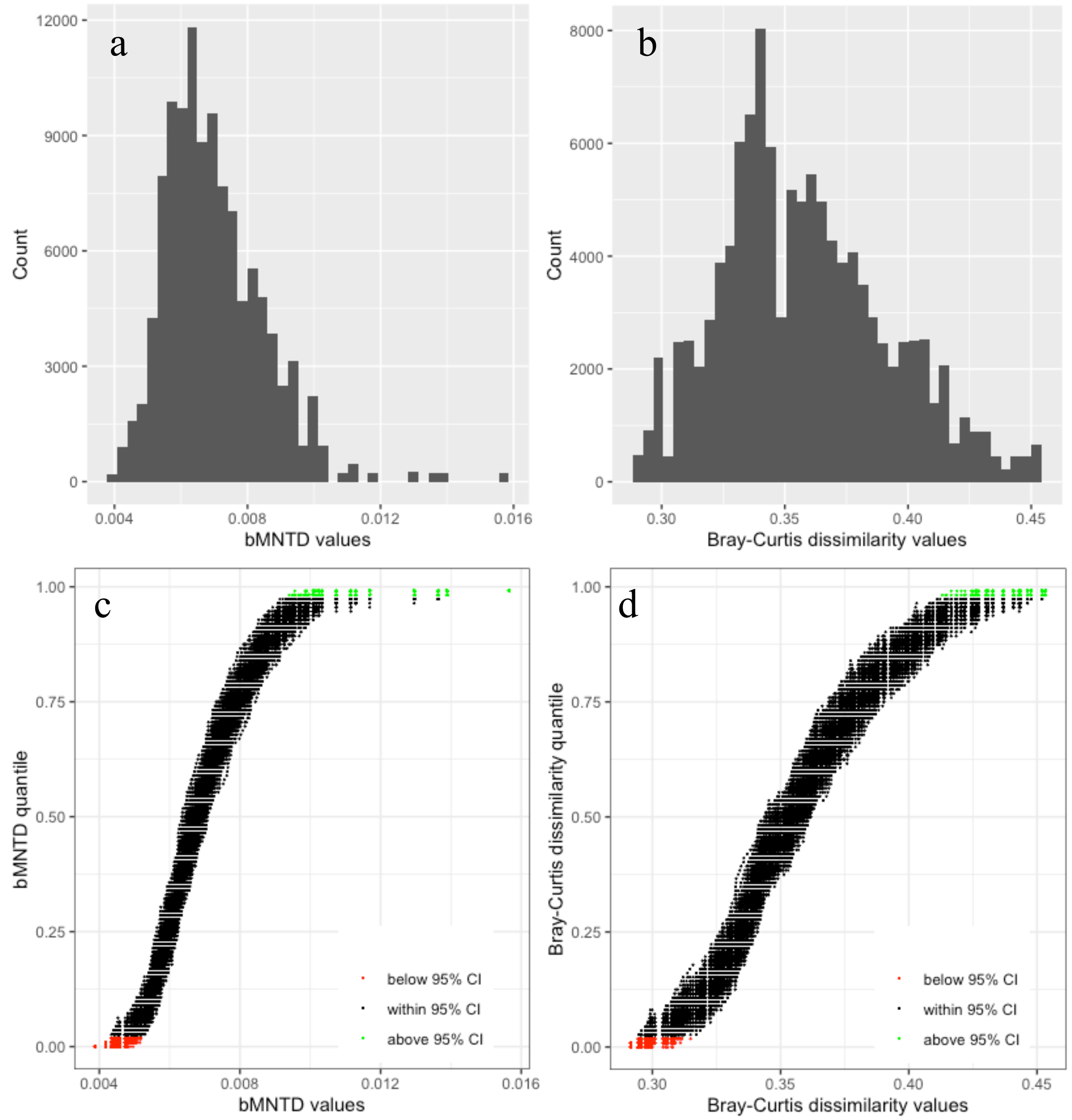

**SI Figure 4**. bMNTD and Bray-Curtis dissimilarity values observed for mixing treatment pairs. Histograms of values for **a**) bMNTD and **b**) Bray-Curtis dissimilarities show the distribution of values across the dataset. Subfigures **c**) and **d**) plot the values as a factor of the quantile in which they fell for each iteration of null model randomization (n = 999). Observed values that fell outside of the central 95% confidence interval are indicated in red (below) and green (above).

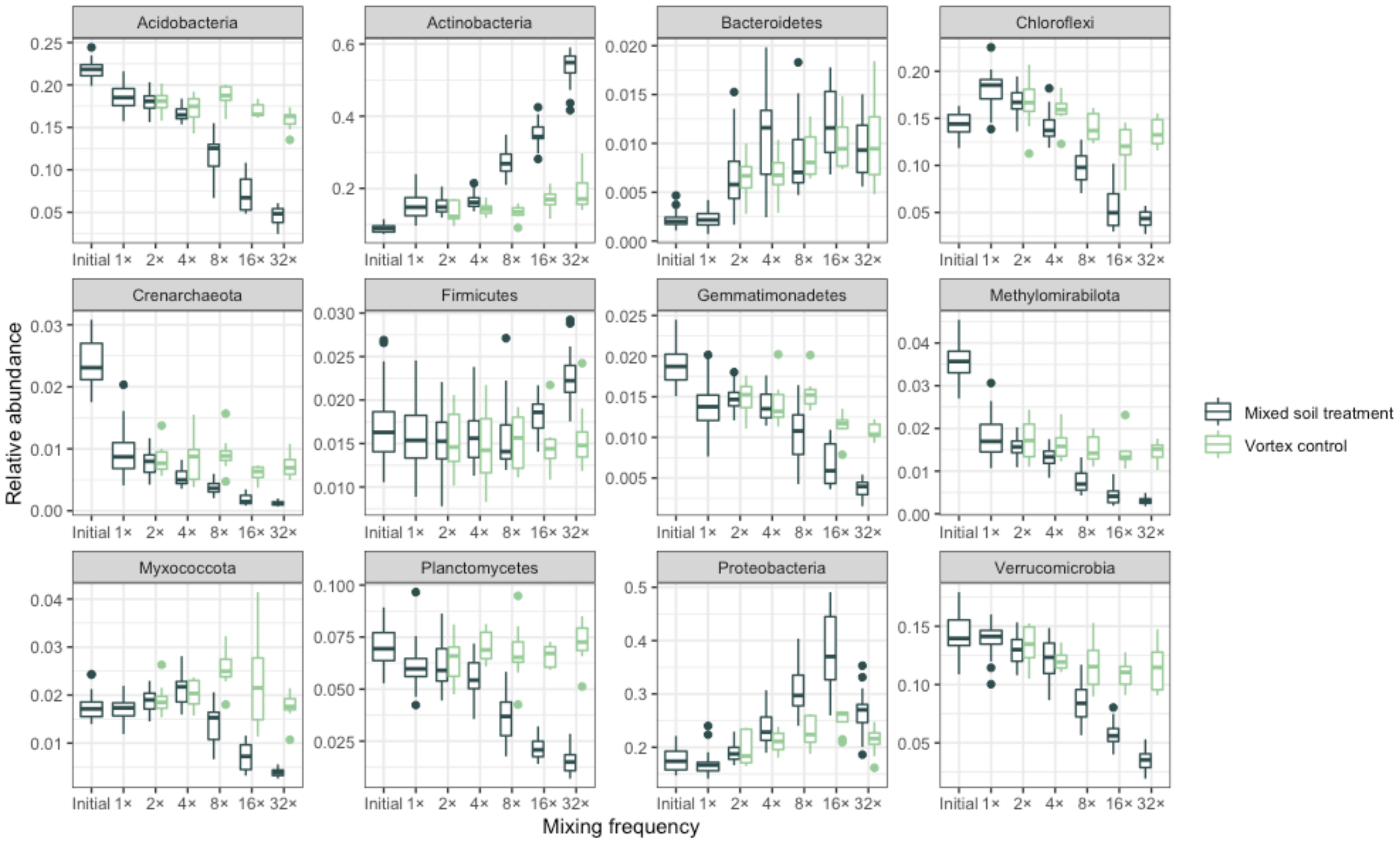

**SI Figure 5**. Relative abundance of the 12 most abundant phyla across initial communities, 1× controls, soil mixing treatments (along x-axis), and vortex controls (by color).

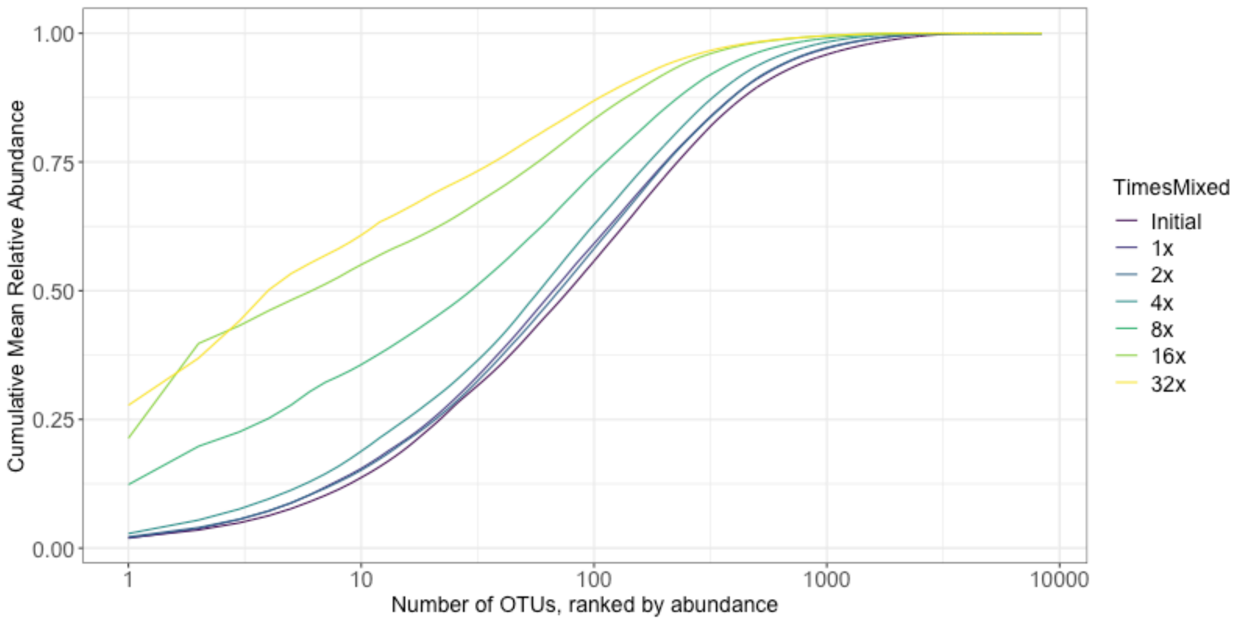

**SI Figure 6**. Cumulative mean relative abundance as a function of abundance-ranked OTUs. At 32×, the single most abundant OTU represents about 30% of mean relative abundance, and the first 10 OTUs comprise over 60% of mean relative abundance. Higher community evenness is apparent in the initial and infrequently mixed soil treatments.

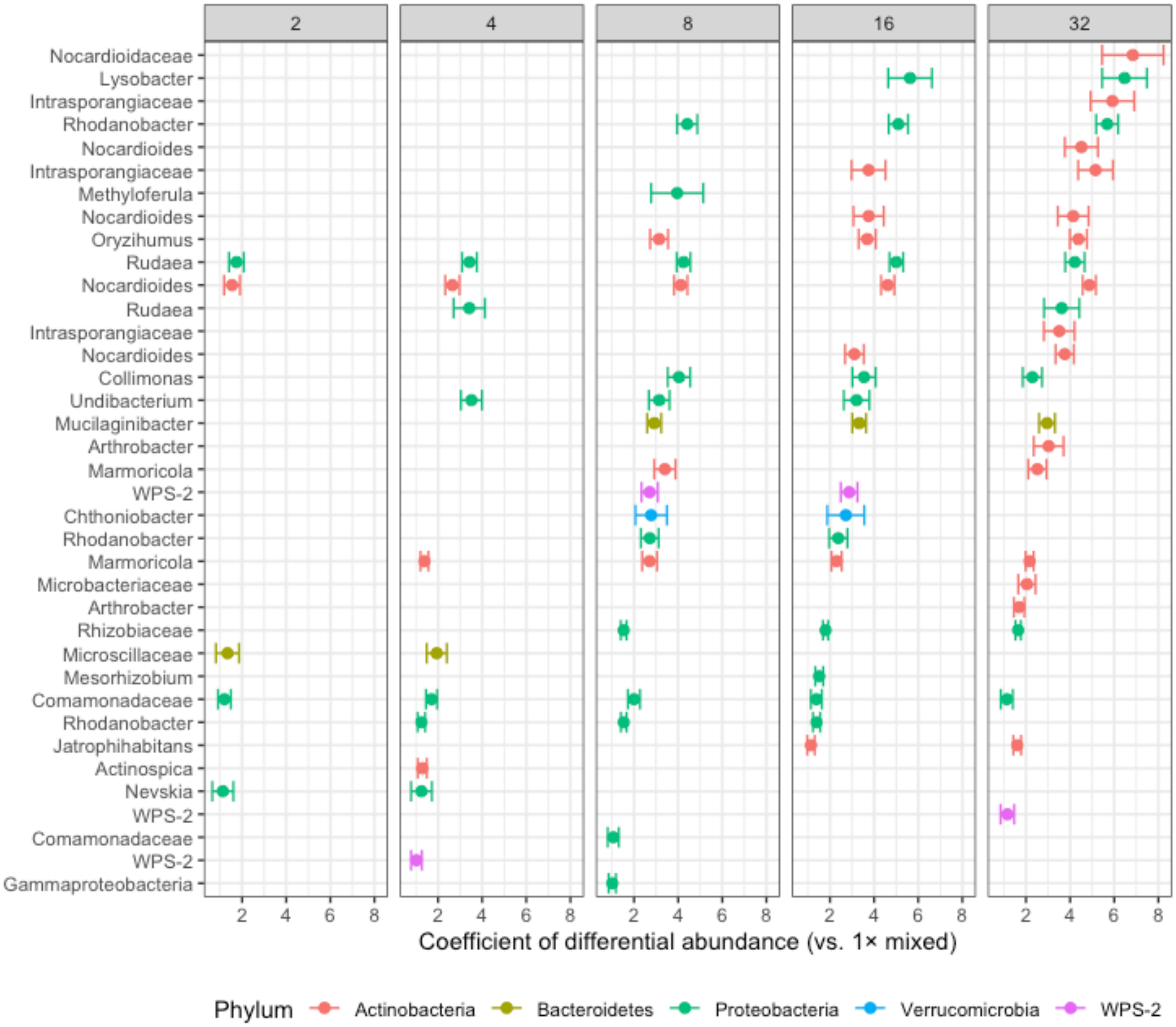

**SI Figure 7**. Bacterial taxa with positive differential abundance (enrichment) in each mixing treatment (excluding vortex controls). The x-axis is the coefficient of differential abundance, in this case (showing enrichment), demonstrating an increase in relative abundance compared to 1×, with only coefficients > 1.0 plotted. Each point represents a single OTU, labelled on the y-axis with the finest available taxonomy and colored by phylum; as such there are several repetitions in name, e.g. *Nocardioides*, as there may have been several OTUs definable to the same name and level of taxonomy.

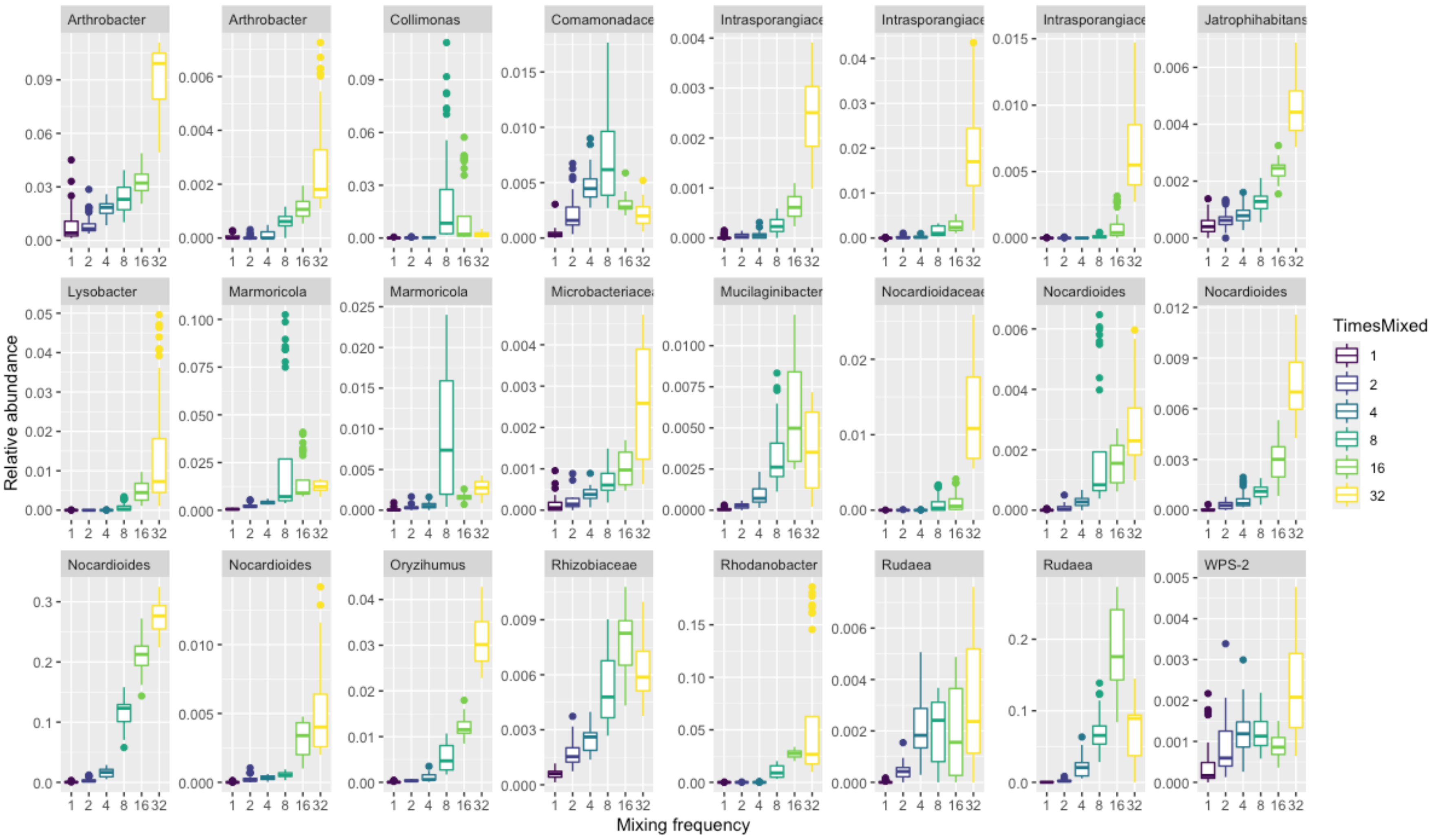

**SI Figure 8**. Bacterial taxa with positive differential abundance (enrichment) at 32× as compared to 1×, presented as relative abundance across all mixing treatments (excluding vortex controls). Facet labels are the finest available taxonomy; as such, there are several repetitions in name, e.g. *Nocardioides*, as there may have been several OTUs definable to the same name and level of taxonomy.

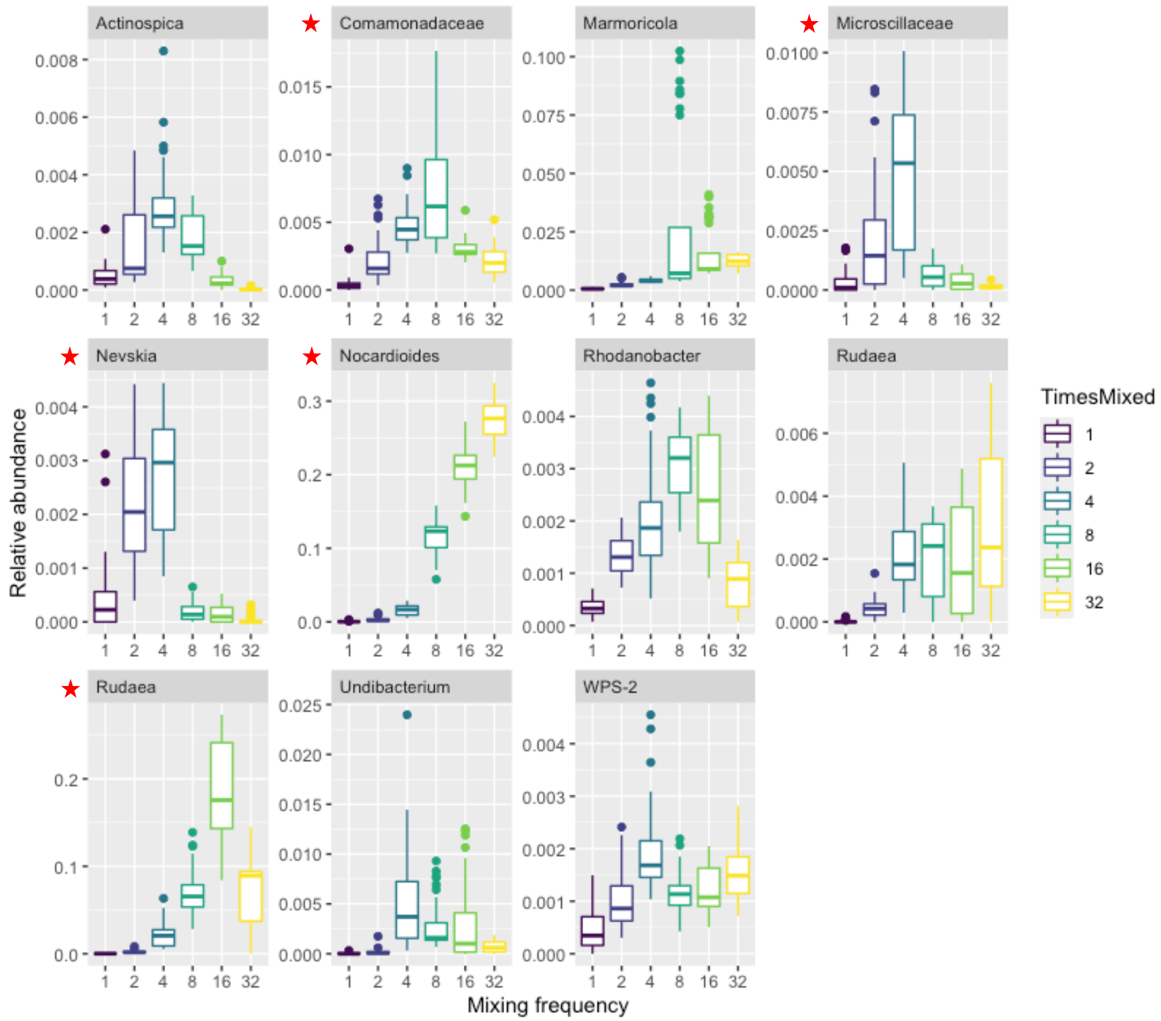

**SI Figure 9**. Bacterial taxa with positive differential abundance (enrichment) at 4× as compared to 1×, presented as relative abundance across all mixing treatments (excluding vortex controls). Red stars mark OTUs that were also significantly enriched at 2×. Facet labels are the finest available taxonomy; as such there may be repetitions in name, e.g. *Rudaea*, as there may have been several OTUs definable to the same name and level of taxonomy.

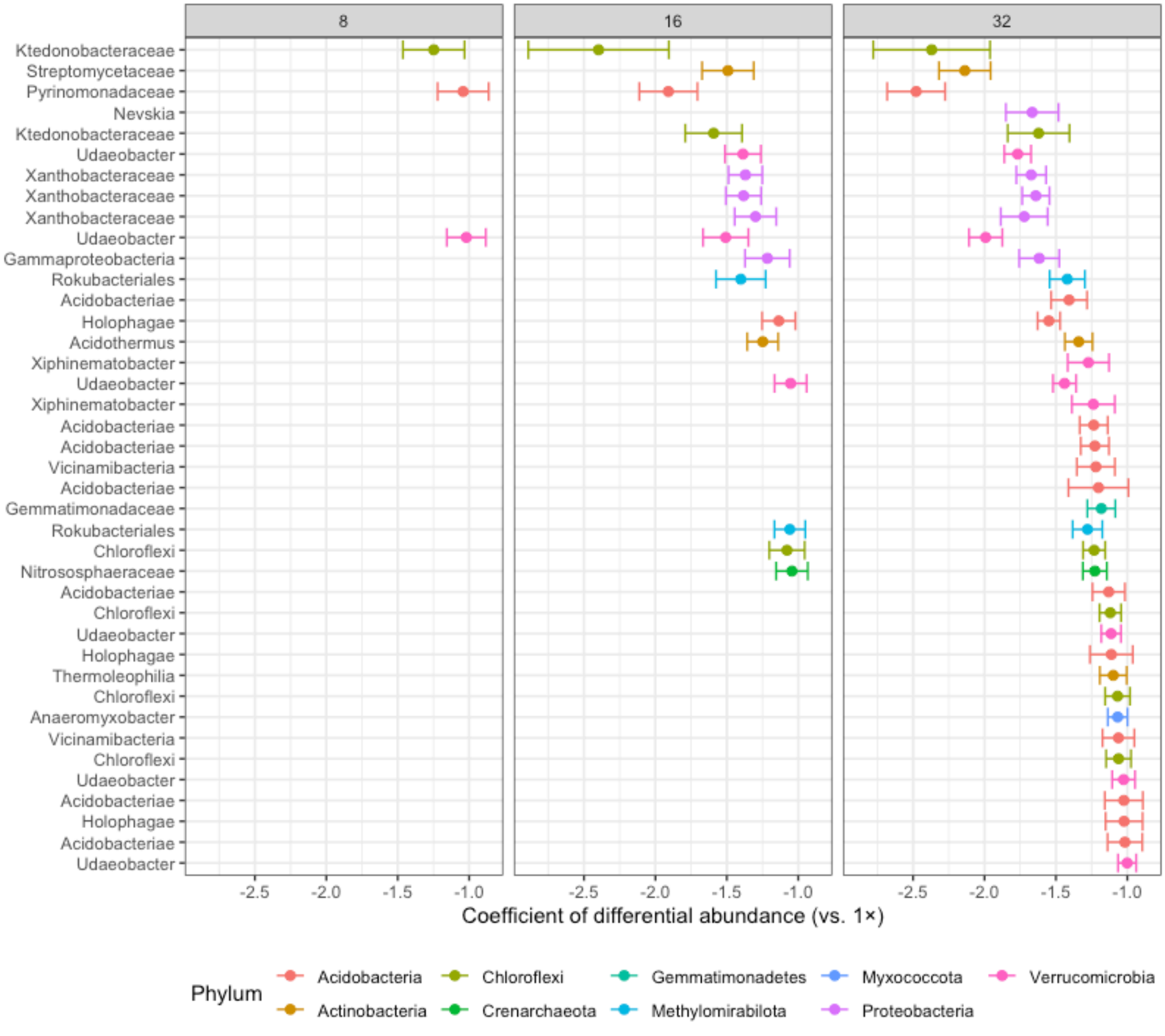

**SI Figure 10**. Bacterial taxa with negative differential abundance (depletion) in each mixing treatment (excluding vortex controls). The x-axis is the coefficient of differential abundance, in this case (showing depletion), demonstrating a decrease in relative abundance compared to 1×, with only coefficients < -1.0 plotted. Each point represents a single OTU, labelled on the y-axis with the finest available taxonomy and colored by phylum; as such there are several repetitions in name, e.g. *Xanthobacteraceae*, as there may have been several OTUs definable to the same name and level of taxonomy.

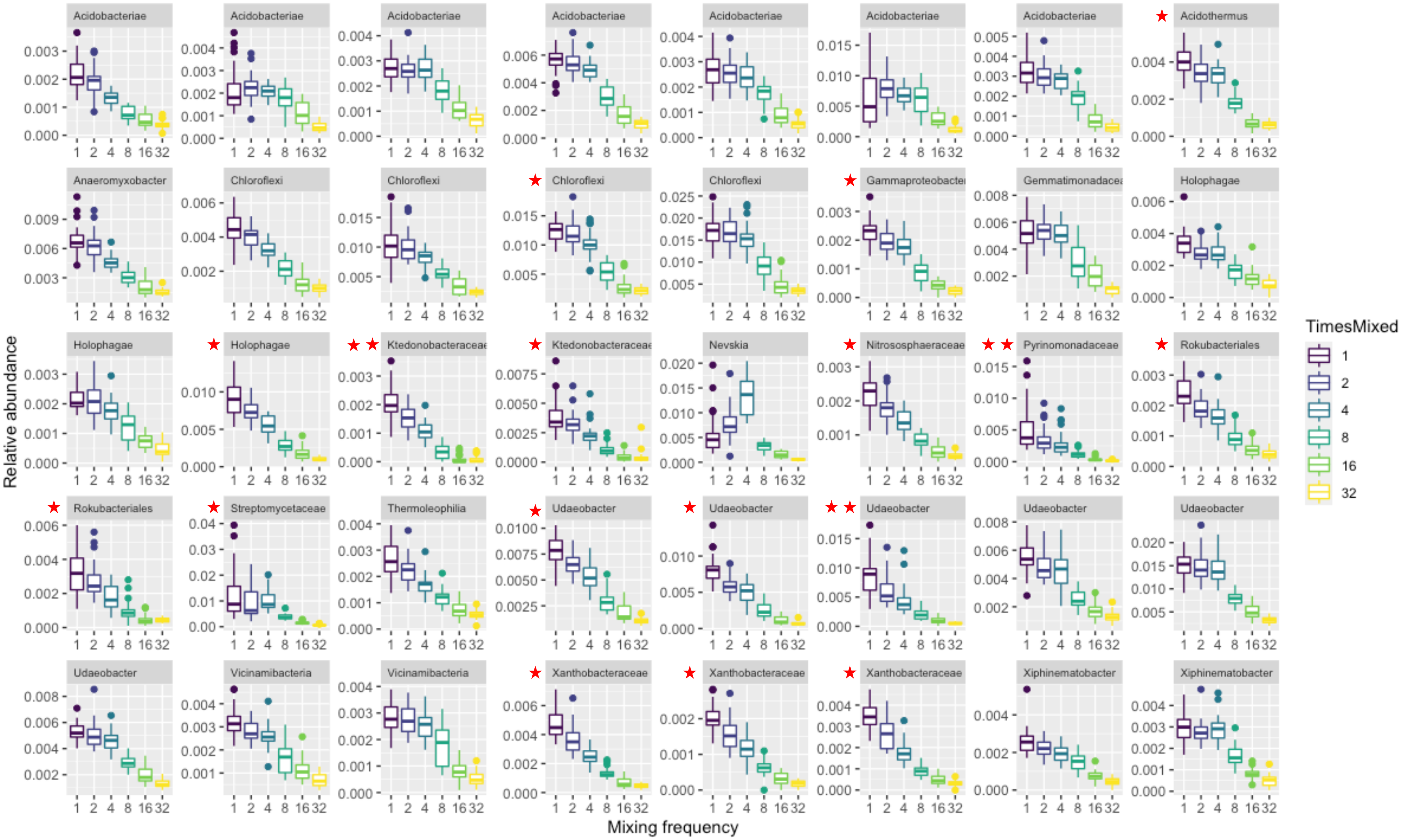

**SI Figure 11**. Bacterial taxa with negative differential abundance (depletion) at 32× as compared to 1×, presented as relative abundance across all mixing treatments (excluding vortex controls). Red stars designate the OTUs that were also depleted at 16× (at least one star) and at 8× (two stars). There were no taxa that were depleted at 4× or 2×, after filtering out very rare taxa. Facet labels are the finest available taxonomy; as such there are several repetitions in name, e.g. *Udaeobacter*, as there may have been several OTUs definable to the same name and level of taxonomy.

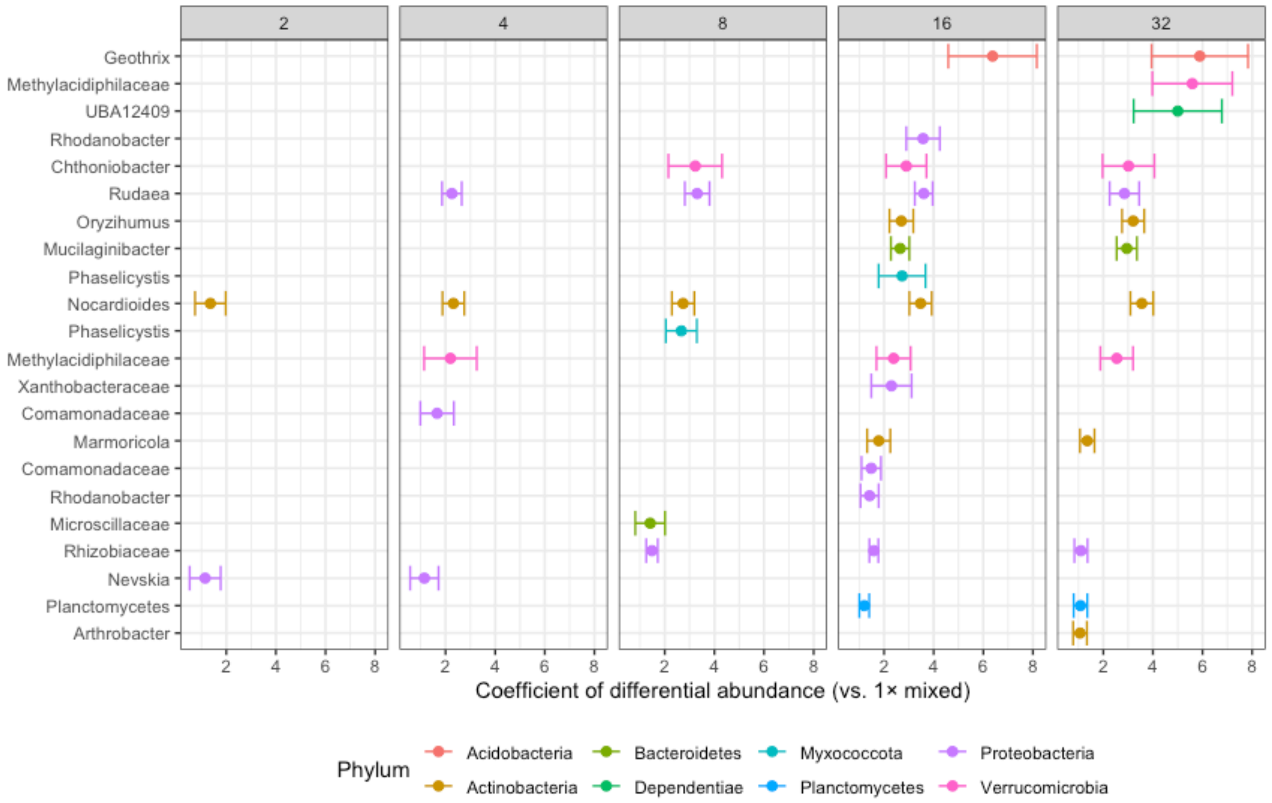

**SI Figure 12**. Bacterial taxa with positive differential abundance (enrichment) in vortex controls. The x-axis is the coefficient of differential abundance, in this case (showing enrichment), demonstrating an increase in relative abundance compared to 1×, with only coefficients > 1.0 plotted. Each point represents a single OTU, labelled on the y-axis with the finest available taxonomy and colored by phylum; as such there are several repetitions in name, e.g. *Rhodanobacter*, as there may have been several OTUs definable to the same name and level of taxonomy.

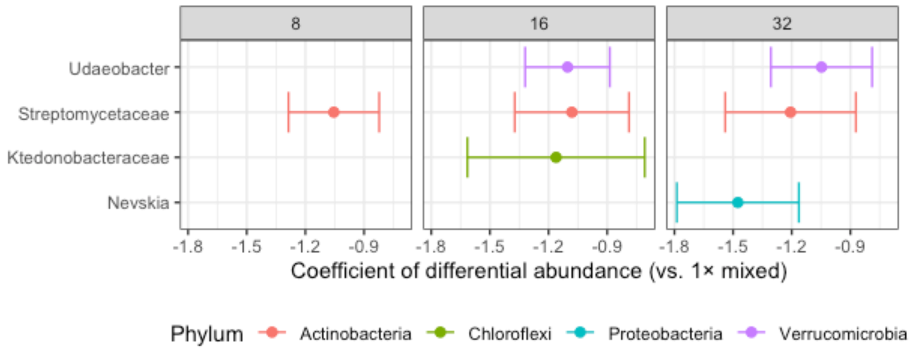

**SI Figure 13**. Bacterial taxa with negative differential abundance (depletion) in vortex controls. The x-axis is the coefficient of differential abundance, in this case (showing depletion), demonstrating a decrease in relative abundance compared to 1×, with only coefficients < -1.0 plotted. Each point represents a single OTU, labelled on the y-axis with the finest available taxonomy and colored by phylum.

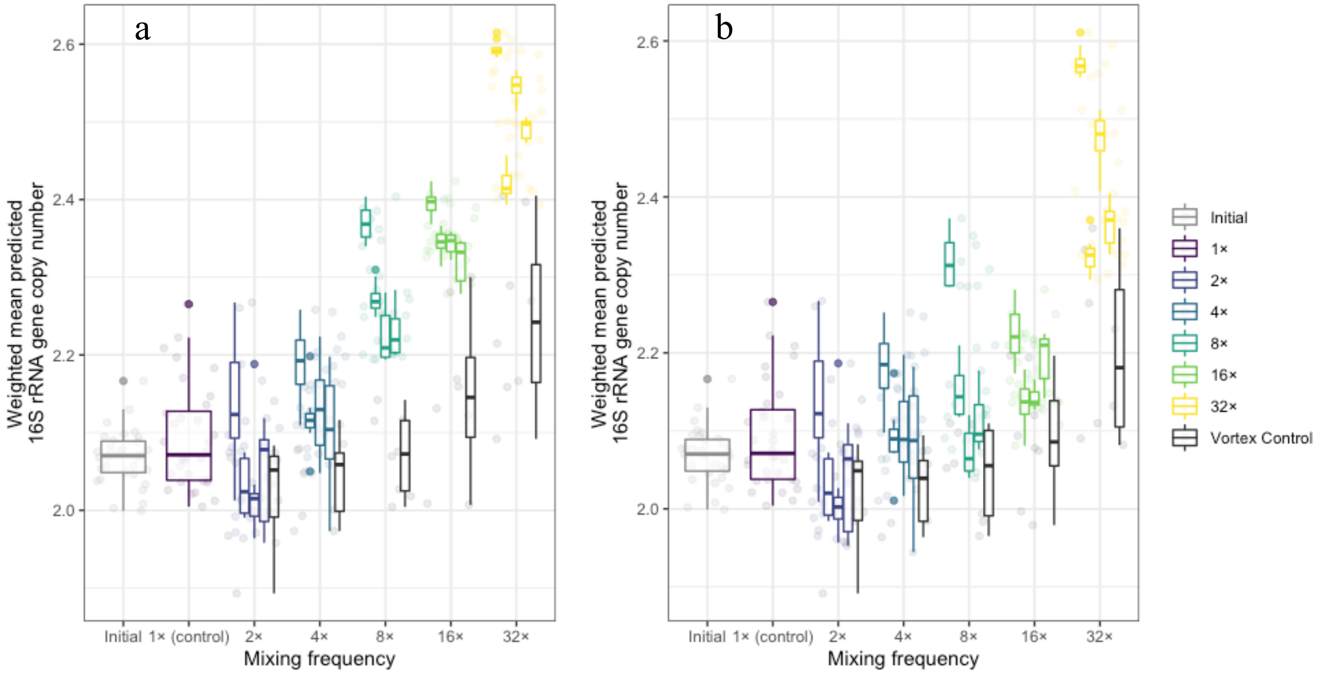

**SI Figure 14**. Weighted mean predicted 16S rRNA gene copy number. These data represent taxa for which a gene copy number was available in the rrnDB (Stoddard et al., 2015) **a**) for all taxa and **b**) excluding OTUs from the genus *Nocardioides*.

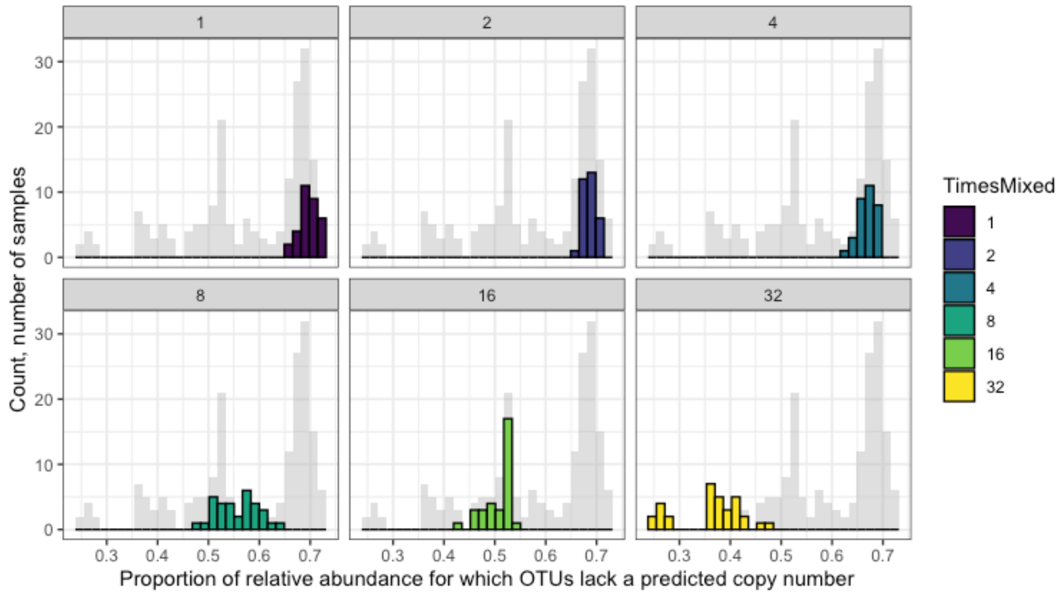

**SI Figure 15**. Histogram of proportion of OTUs with no copy number available in rrnDB. The bimodal aspect of the overall dataset (in gray) is driven by the dominance of the genus *Nocardioides* in the more frequently mixed treatments, for which a predicted copy number was available, thus decreasing the proportion of total relative abundance lacking a predicted copy number.

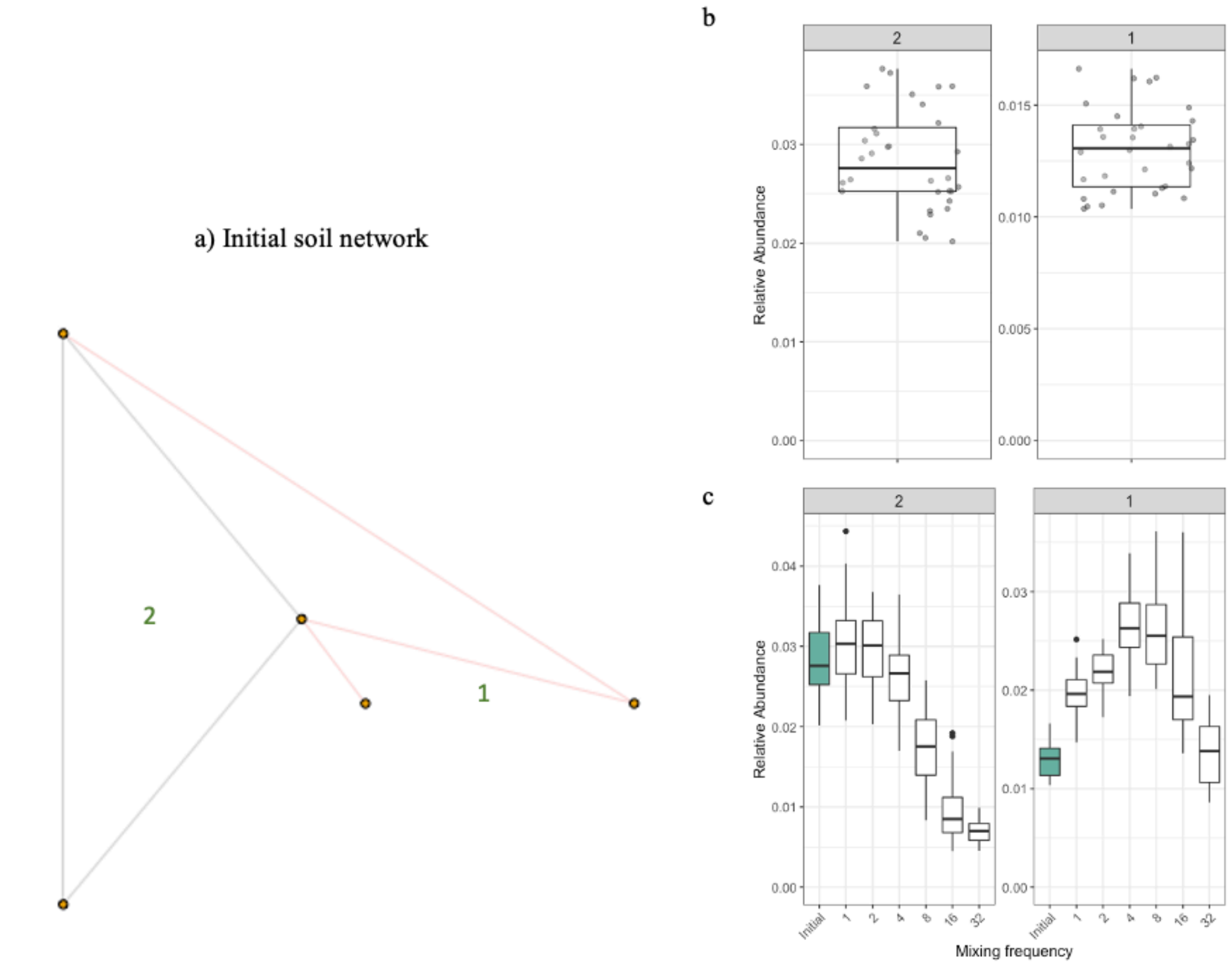

**SI Figure 16**. Co-occurrence network, Initial samples. **a**) Network constructed from initial soil samples that did not undergo incubation. Edges (lines) connecting nodes (points; OTUs) that co-occur are in black, and edges connecting nodes that represent co-exclusions are in red. Module numbers (green) are arbitrary and correspond to the facet titles in the boxplot figures. **b**) Relative abundance of OTUs within each module, plotted for each sample (n=32). **c**) Relative abundance of OTUs within each module, by mixing treatment; the initial treatment is filled in. The sparsity and arguably uninformative nature of this network underscores the stochasticity of these random subsamples of homogenized soil (Faust, 2021). Note that module numbers are assigned arbitrarily for reference and are not consistent across networks for different mixing treatments.

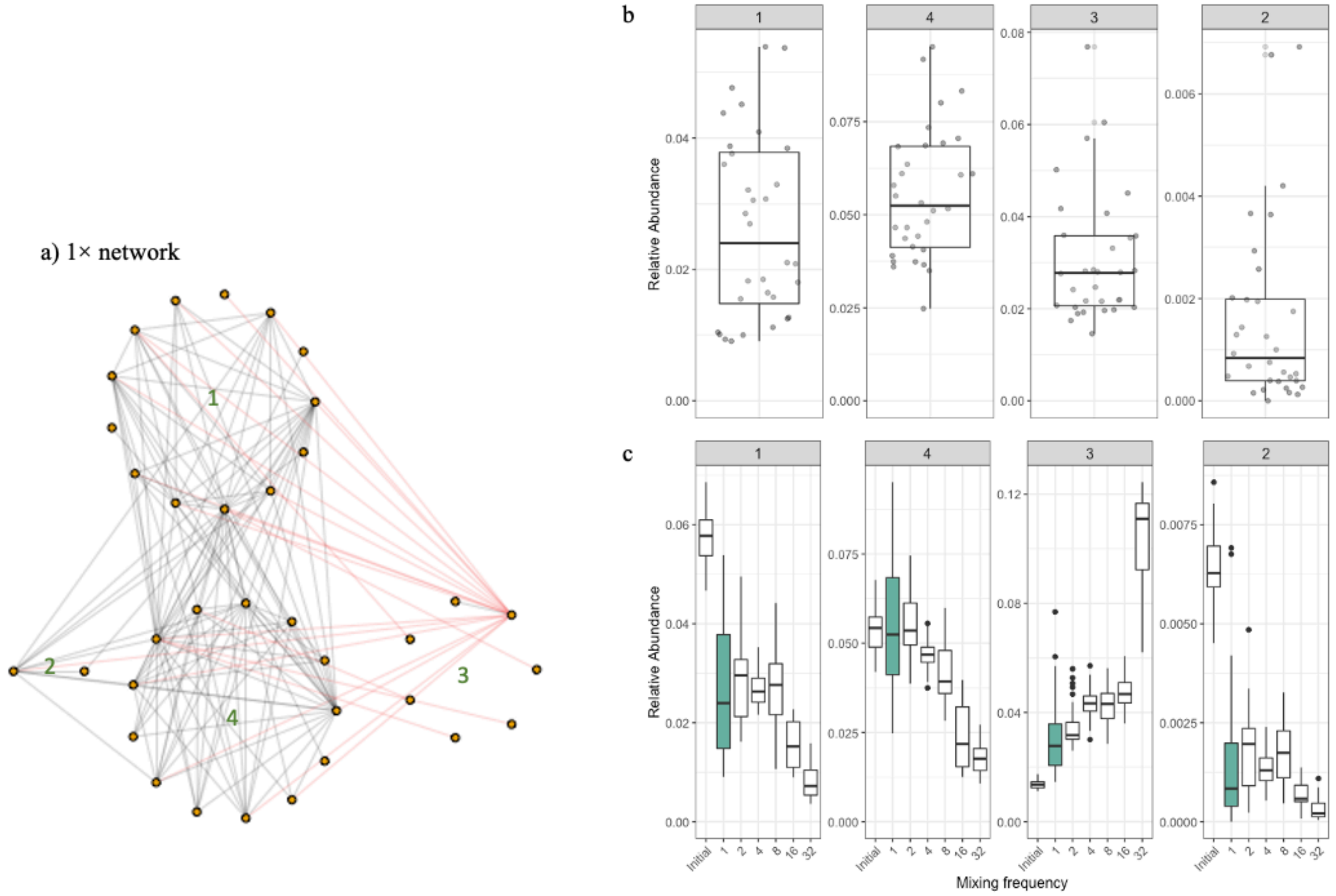

**SI Figure 17**. Co-occurrence network, 1× samples. **A**) Network constructed from 1× control soil. Edges (lines) connecting nodes (points; OTUs) that co-occur are in black, and edges connecting nodes that represent co-exclusions are in red. Module numbers (green) are arbitrary and correspond to the facet titles in the boxplot figures. **B**) Relative abundance of OTUs within each module, plotted for each 1× sample (n=32). **C**) Relative abundance of OTUs within each module, by mixing treatment; the 1× treatment is filled in. Note that module numbers are assigned arbitrarily for reference and are not consistent across networks for different mixing treatments.

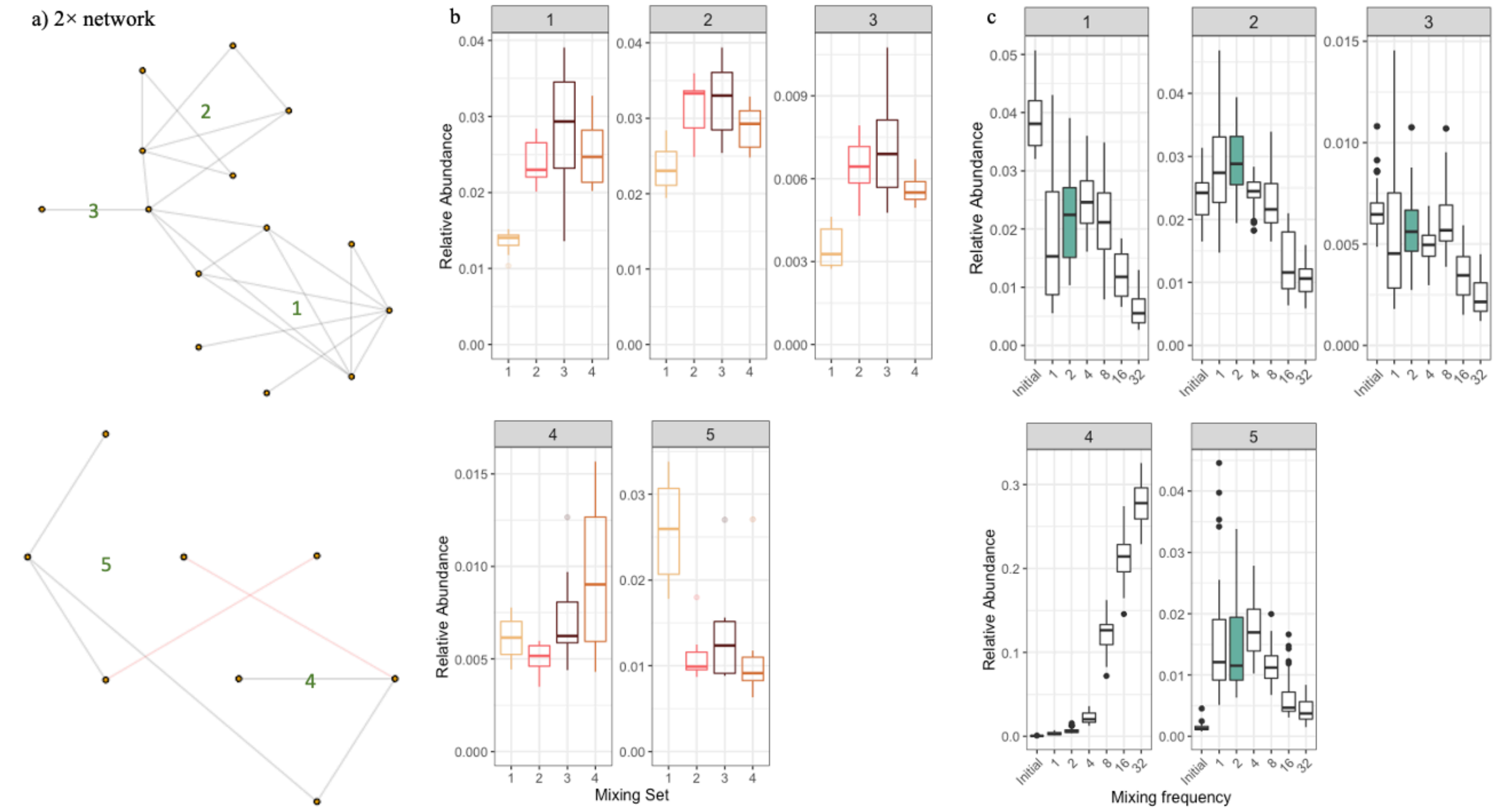

**SI Figure 18**. Co-occurrence network, 2× treatment samples. **A**) Network for the 2× treatment, with a main component (modules 1 – 3) and a secondary component (modules 4 & 5). Edges (lines) connecting nodes (points; OTUs) that co-occur are in black, and edges connecting nodes that represent co-exclusions are in red. Module numbers (green) are arbitrary and correspond to the facet titles in the boxplot figures. **B**) Relative abundance of OTUs within each module, by mixing set. **c**) Relative abundance of OTUs within each module, by mixing treatment; the 2× treatment is filled in. Note that module numbers are assigned arbitrarily for reference and are not consistent across networks for different mixing treatments, and that mixing sets are not consistent across treatments (e.g. mixing set #2 at 2× is totally unrelated to mixing set #2 at 8×).

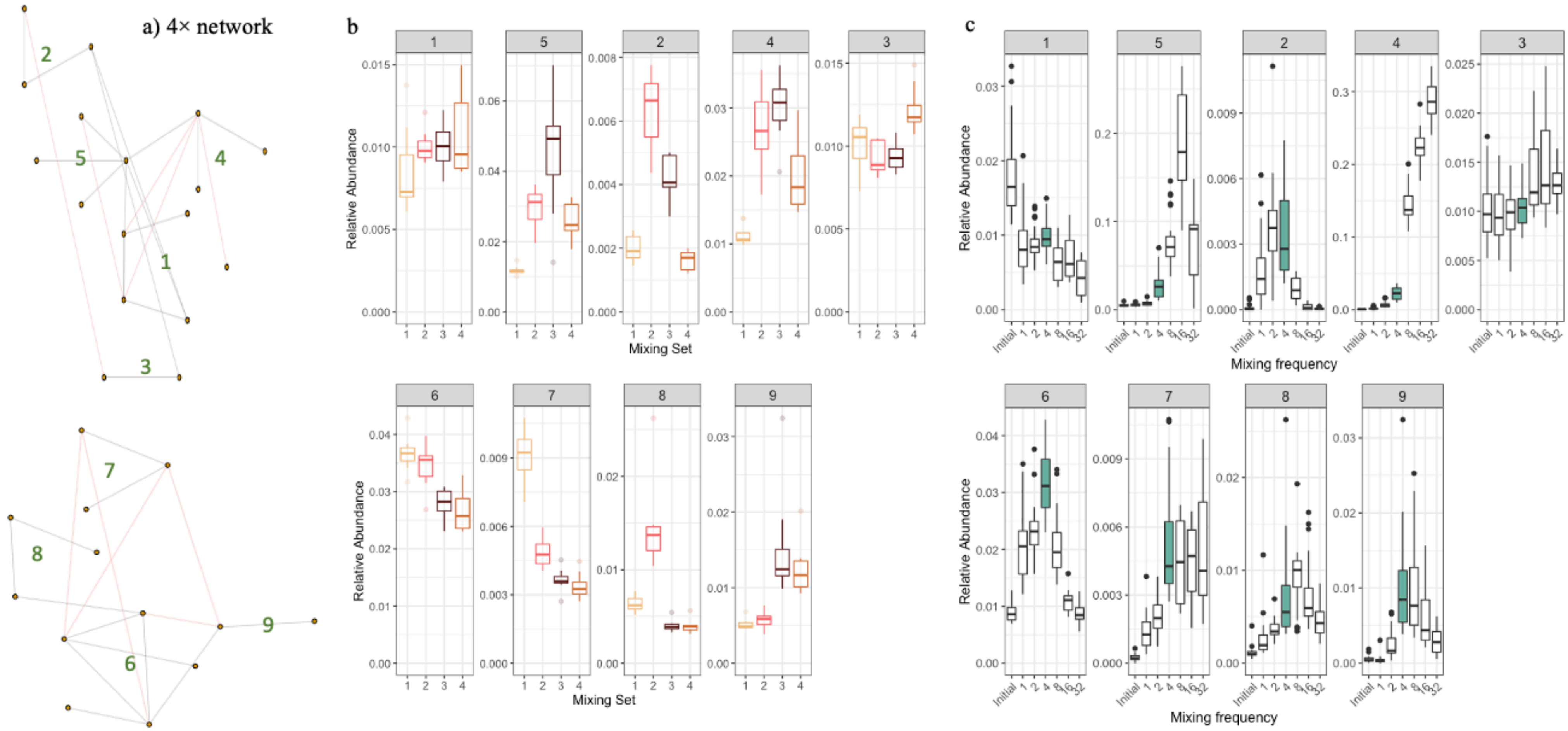

**SI Figure 19**. Co-occurrence network, 4× treatment samples. **a**) Network for the 4× treatment, with a main component (modules 1 - 5) and secondary component (modules 6 - 9). Edges (lines) connecting nodes (points; OTUs) that co-occur are in black, and edges connecting nodes that represent co-exclusions are in red. Module numbers (green) are arbitrary and correspond to the facet titles in the boxplot figures. **b**) Relative abundance of OTUs within each module, by mixing set. **c**) Relative abundance of OTUs within each module, by mixing treatment; the 4× treatment is filled in. Note that module numbers are assigned arbitrarily for reference and are not consistent across networks for different mixing treatments, and that mixing sets are not consistent across treatments (e.g. mixing set #2 at 2× is totally unrelated to mixing set #2 at 8×).

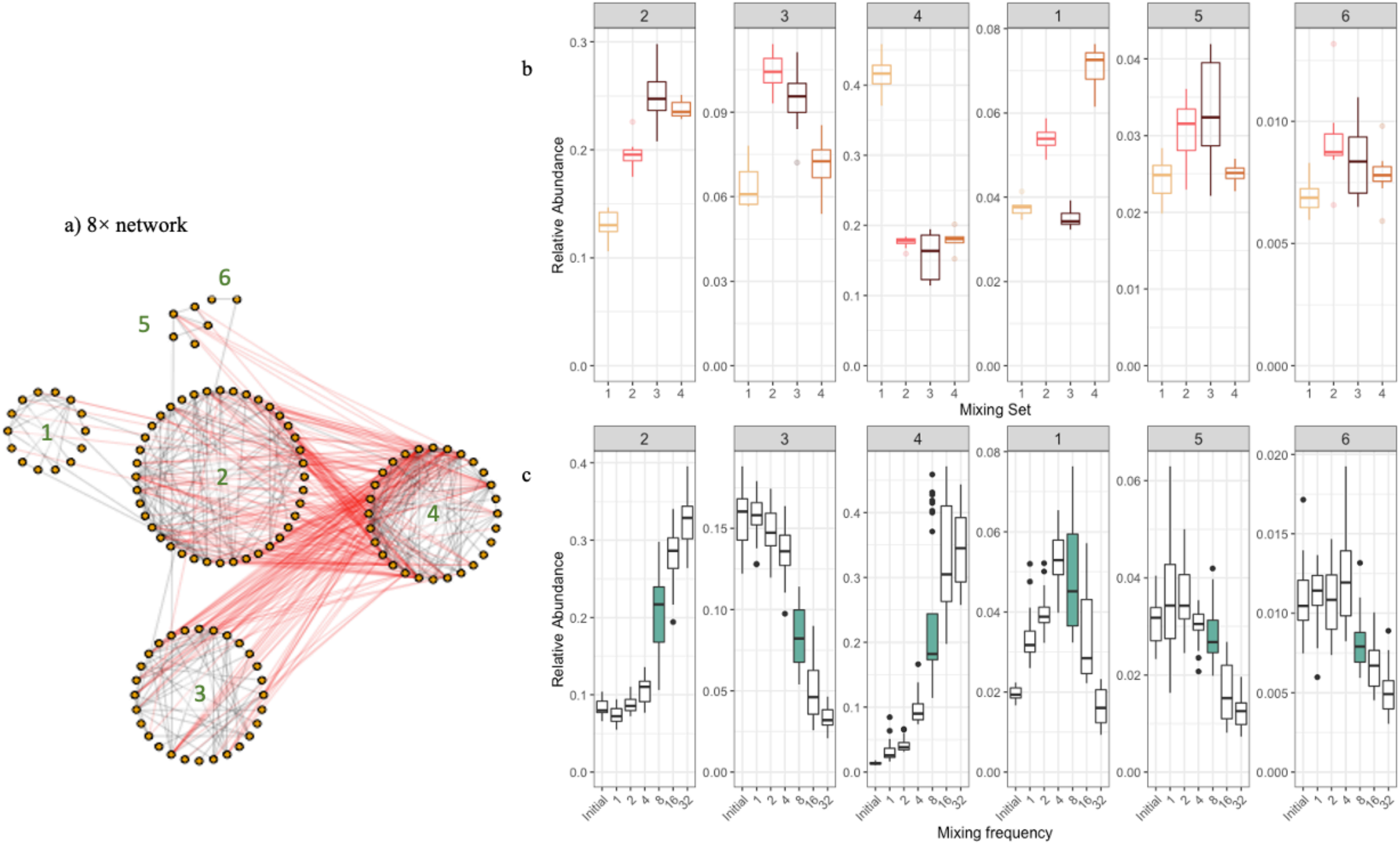

**SI Figure 20**. Co-occurrence network, 8× treatment samples. **a**) Network for the 8× treatment. Edges (lines) connecting nodes (points; OTUs) that co-occur are in black, and edges connecting nodes that represent co-exclusions are in red. Module numbers (green) are arbitrary and correspond to the facet titles in the boxplot figures. **b**) Relative abundance of OTUs within each module, by mixing set. **c**) Relative abundance of OTUs within each module, by mixing treatment; the 8× treatment is filled in. Note that module numbers are assigned arbitrarily for reference and are not consistent across networks for different mixing treatments, and that mixing sets are not consistent across treatments (e.g. mixing set #2 at 2× is totally unrelated to mixing set #2 at 8×).

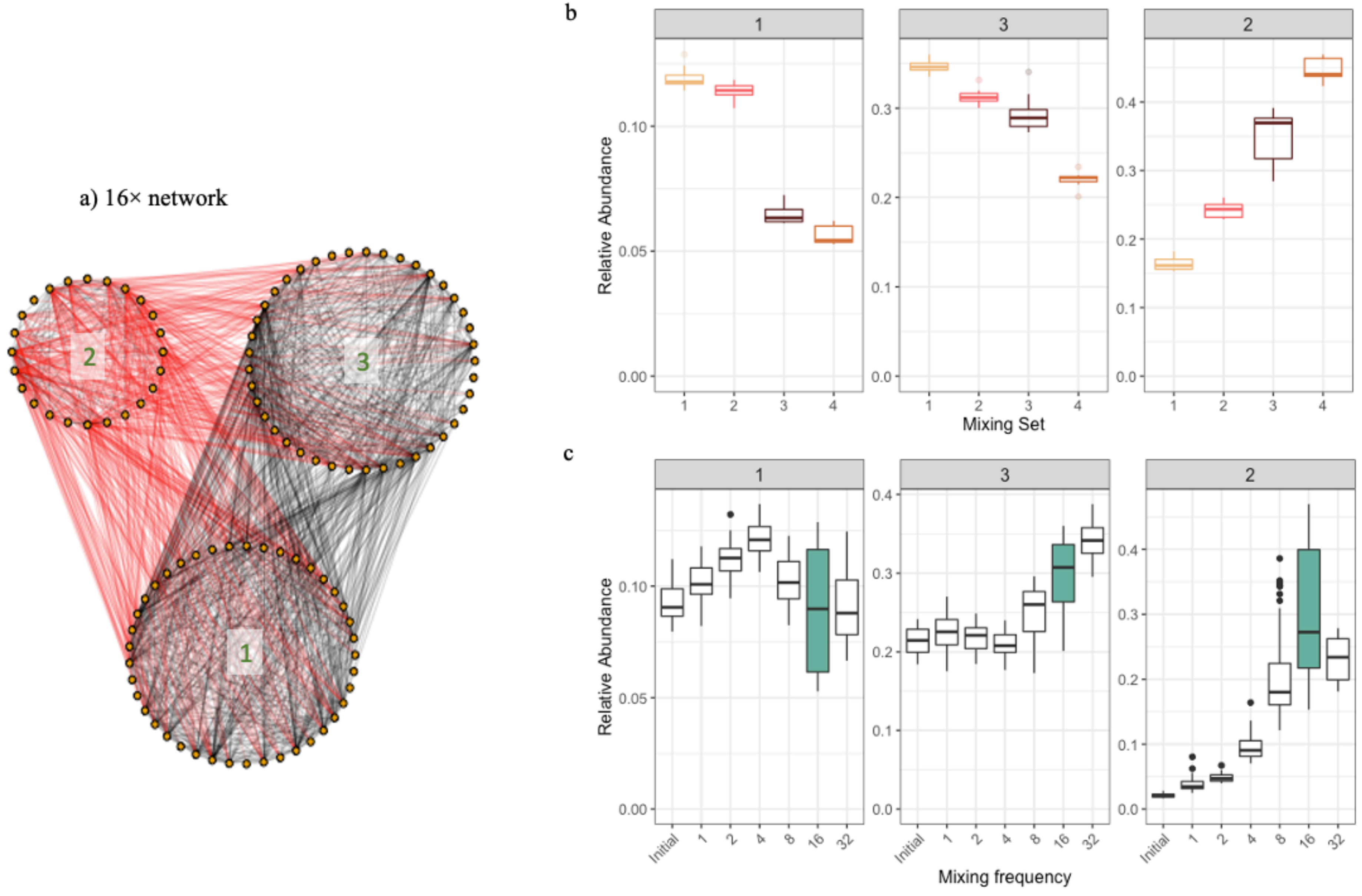

**SI Figure 21**. Co-occurrence network, 16× treatment samples. **a**) Network for the 16× treatment. Edges (lines) connecting nodes (points; OTUs) that co-occur are in black, and edges connecting nodes that represent co-exclusions are in red. Module numbers (green) are arbitrary and correspond to the facet titles in the boxplot figures. **b**) Relative abundance of OTUs within each module, by mixing set. **c**) Relative abundance of OTUs within each module, by mixing treatment; the 16× treatment is filled in. Note that module numbers are assigned arbitrarily for reference and are not consistent across networks for different mixing treatments, and that mixing sets are not consistent across treatments (e.g. mixing set #2 at 2× is totally unrelated to mixing set #2 at 8×).

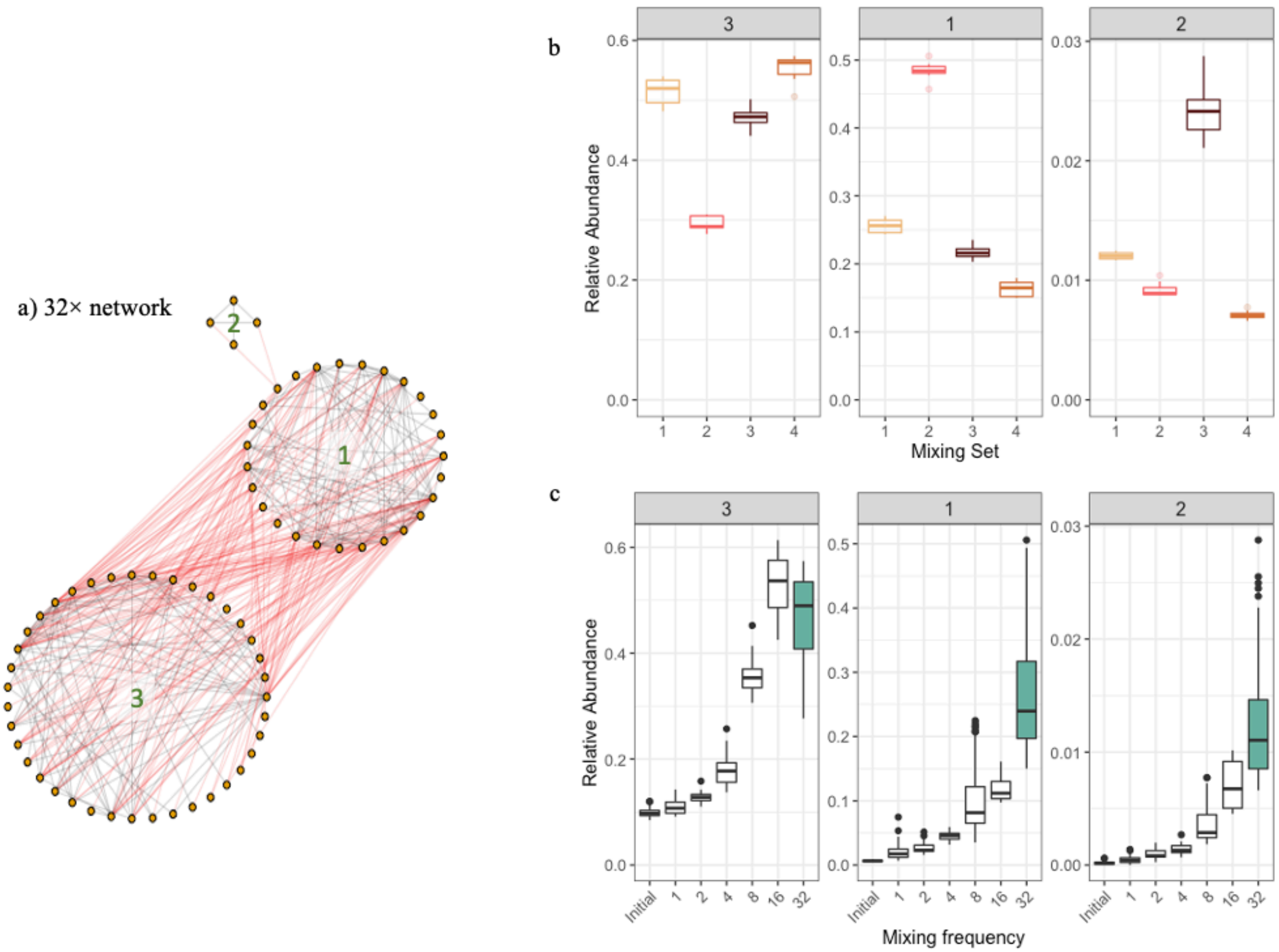

**SI Figure 22**. Co-occurrence network, 32× treatment samples. **a**) Network for the 32× treatment. Edges (lines) connecting nodes (points; OTUs) that co-occur are in black, and edges connecting nodes that represent co-exclusions are in red. Module numbers (green) are arbitrary and correspond to the facet titles in the boxplot figures. **b**) Relative abundance of OTUs within each module, by mixing set. **c**) Relative abundance of OTUs within each module, by mixing treatment; the 32× treatment is filled in. Note that module numbers are assigned arbitrarily for reference and are not consistent across networks for different mixing treatments, and that mixing sets are not consistent across treatments (e.g. mixing set #2 at 2× is totally unrelated to mixing set #2 at 8×).

#### SI References

Faust, K., 2021. Open challenges for microbial network construction and analysis. The ISME Journal 1–8. doi:10.1038/s41396-021-01027-4

Kozich, J.J., Westcott, S.L., Baxter, N.T., Highlander, S.K., Schloss, P.D., 2013. Development of a Dual-Index Sequencing Strategy and Curation Pipeline for Analyzing Amplicon Sequence Data on the MiSeq Illumina Sequencing Platform. Appl. Environ. Microbiol. 79, 5112–5120. doi:10.1128/aem.01043-13

Stoddard, S.F., Smith, B.J., Hein, R., Roller, B.R.K., Schmidt, T.M., 2015. rrnDB: improved tools for interpreting rRNA gene abundance in bacteria and archaea and a new foundation for future development. Nucleic Acids Research 43, D593–D598. doi:10.1093/nar/gku1201

Walters, W., Hyde, E.R., Berg-Lyons, D., Ackermann, G., Humphrey, G., Parada, A., Gilbert, J.A., Jansson, J.K., Caporaso, J.G., Fuhrman, J.A., Apprill, A., Knight, R., Bik, H., 2016. Improved Bacterial 16S rRNA Gene (V4 and V4-5) and Fungal Internal Transcribed Spacer Marker Gene Primers for Microbial Community Surveys. MSystems 1, e00009-15. doi:10.1128/msystems.00009-15
